# Coupled Binding and Folding of NS2B/NS3 Protease and Linker Effects Revealed by Topology-based Modeling

**DOI:** 10.64898/2026.05.04.722635

**Authors:** Kairong Dong, Jian Huang, Min Chen, Jianhan Chen

## Abstract

Orthoflavivirus, such as West Nile Virus (WNV), dengue virus (DENV) and ZIKA virus (ZIKV), are globally distributed pathogens that pose substantial threats to human health. Currently, there are still no effective antiviral drugs for WNV or ZIKV. Despite the availability of two licensed DENV vaccines, their use remains limited due to potential risks, highlighting an urgent need for antiviral drug development. The highly conserved orthoflavivirus protease NS2B/NS3 is required for viral replication, making it a promising anti-flavivirus target. A major challenge, however, is the highly charged active site of this enzyme, which requires charged chemical matters with low bioavailability. An alternative and more attractive strategy is to target potential allosteric sites or folding intermediate states of the protease. In this work, we employ the topology-based coarse-grained Gō modeling to explore the coupled binding and folding pathways of WNV NS2B/NS3 protease and study the effects of the widely used experimental construct with a G4SG4 linker between NS2B and NS3 on stability and folding. Our results provide a holistic conformational landscape of the protease binding and folding, including several key intermediate states. We find that the presence of the G4SG4 linker alters the folding pathways and destabilizes the NS2B C-terminus. The latter is consistent with experimental observations that the G4SG4 linked protease has lower activity and adopts an open state without the substrate in crystal structures. Together, these findings provide for the first time a complete picture of the binding and folding of the NS2B/NS3 protease and identify important folding intermediate states that could be targeted for allosteric antiviral drug development.

**TOC Figure:** 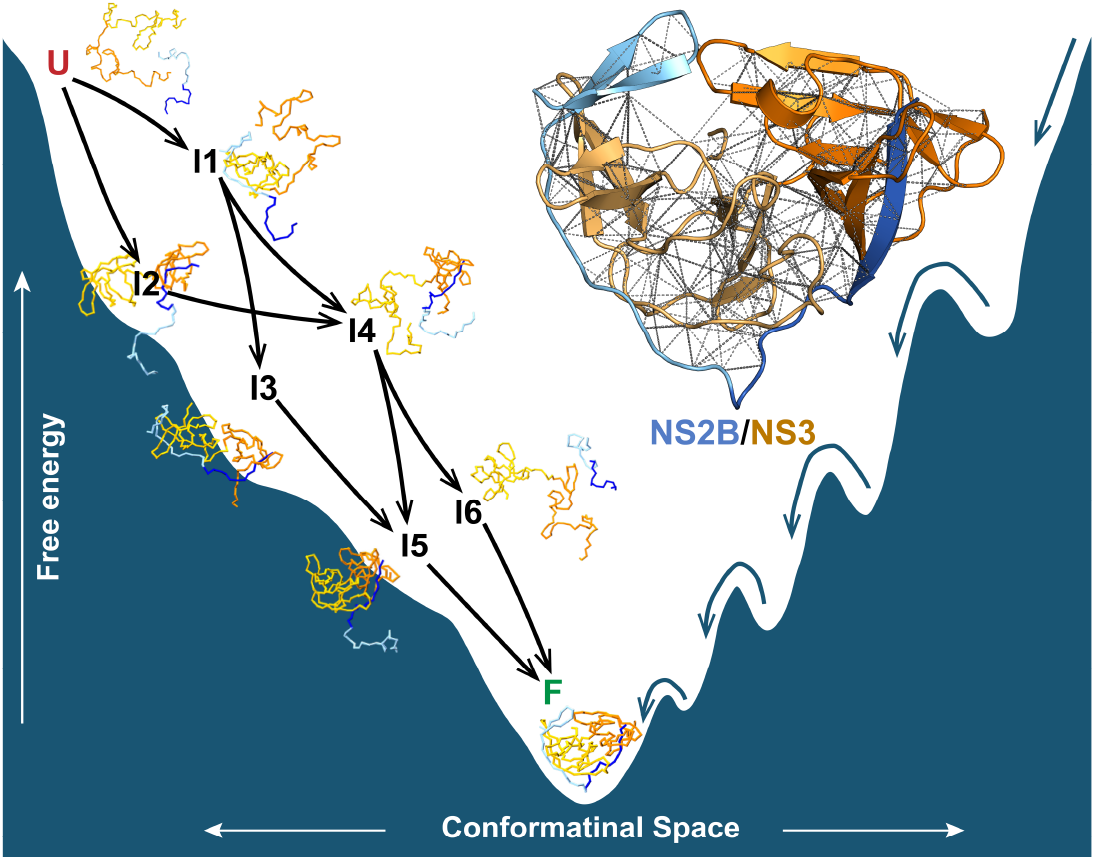

## Introduction

Orthoflavivirus is a genus of arthropod-borne viruses that includes dengue virus (DENV), West Nile virus (WNV), Zika virus (ZIKV), tick-borne encephalitis virus (TBEV), yellow fever virus (YFV), and others ^1^. Among orthoflavivirus members, the mosquito-borne DENV, WNV, and ZIKV can cause many life-threatening diseases, including vascular shock syndrome and encephalitis ^2^. Additionally, ZIKV is known to cause congenital abnormalities and fetal death ^3^. Over the past decades, Orthoflavivirus has spread dramatically worldwide. DENV has infected over 3 billion people in endemic regions and now infects up to 400 million people annually ^4,5^; WNV has rapidly expanded in the United States and other regions ^4^. Unfortunately, effective antiviral treatments are still sparse for flavivirus infections. Currently, except for YFV, vaccine development for most orthoflavivirus has faced significant challenges ^6–10^, resulting in no licensed vaccines for the wide-spreading WNV and ZIKA. Even though two licensed DENV vaccines DengvaxiaⓇ/CYD-TDV by Sanofi and QdengaⓇ/TAK003 by Takeda are available ^11–13^, their applications are constrained by age range or pre-vaccination screening due to the risk of causing the severe dengue fever and dengue hemorrhagic fever ^14^. Therefore, there is an unmet need for anti-virial drug development, which likely requires a deeper molecular-level understanding of the virial infection and progression process.

Orthoflavivirus are small-sized enveloped viruses encapsulating a single positive-sense RNA genome that encodes a single polyprotein chain. During the replication phase of viral infection, the polyprotein chain needs to be cleaved by host cell proteases and the viral protease into three structural proteins (capsid, pre-membrane and envelop proteins) and seven non-structural proteins (NS1, NS2A, NS2B, NS3, NS4A, NS4B and NS5)^15,16^. Among the non-structural proteins, the NS5 polymerase and the NS2B/NS3 protease are attractive drug targets for their essential roles in viral cell cycle, high conservation among orthoflaviviruses ^17–21^. However, development of NS5 inhibitor drugs has been hindered by poor bioactivation of candidate compounds and toxicity ^22,23^. Inhibitors of the NS2B/NS3 protease have been successfully developed for hepatitis C virus (HCV) and human immunodeficiency virus (HIV), highlighting its great potential as a target for anti-orthoflavivirus drug design ^22,24,25^.

Orthoflavivirus protease is a serine protease containing two segments, the NS3 protease core and the intrinsically disordered NS2B cofactor ^26,27^. The whole NS3 is a multidomain protein that contains an N-terminal serine protease core domain (∼180 amino acid long) and a C-terminal helicase domain^25^. For simplicity, we refer to the NS3 protease core domain hereafter as NS3. A fragment of ∼40 residues from the NS2B hydrophilic part is necessary and sufficient to form an enzymatically active complex with NS3 ^28–30^. Several structures of the NS2B/NS3 proteases have been determined for WNV, ZIKV and DENV with or without inhibitors ^18,31–36^. Overall, those structures can be categorized into two states: the closed active state and the open non-active state ^31–38^. In both states, the NS3 domain consists of two β-barrels, denoted the N-and C-terminal barrels (NS3N and NS3C), respectively (Fig. 1 A & B).

**Figure 1.**
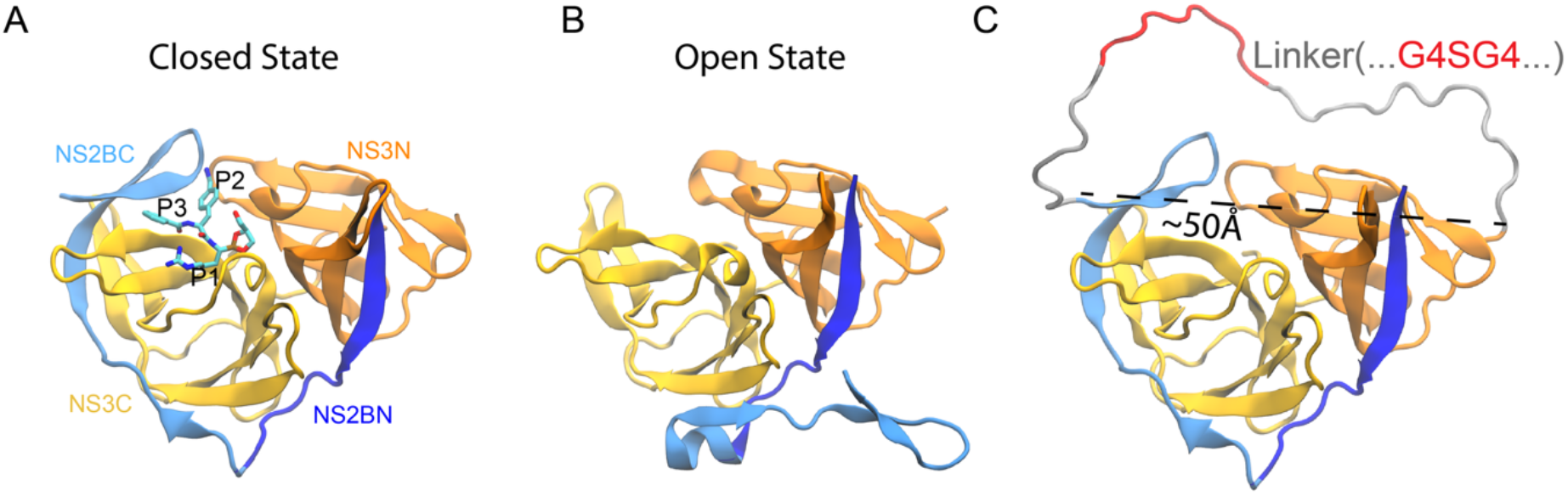
Representative conformational states of WNV NS2B/NS3 protease complexes. **A)** A ligand-bound closed active state (PDB 5IDK ^42^); **B)** An open non-active state (PDB 2GGV ^43^). The N-terminal and C-terminal segments of NS2B, NS2BN and NS2BC domains, are colored deep and light blue; The two barrels, NS3N and NS3C domains, are colored orange and yellow respectively. The bound inhibitor in the closed state is shown using sticks with C, N, O atoms colored cyan, blue and red respectively. **C)** The protease in the closed state with the G4SG4 linker (the “linked” construct) was generated using the AlphaFold2 colab (see Methods). The G4SG4 linker and the disordered residues of the protease are colored red and gray, respectively.

The NS3N barrel includes a strand from the NS2B N-terminal segment (NS2BN), while NS3C interacts with the C-terminal hairpin of NS2B (NS2BC). The main structural difference between the open and closed state lies in the conformation of NS2BC: in the closed active state, the NS2BC wraps around the NS3C barrel and locates in vicinity to the activity site (Fig. 1A); while in the open state, the NS2BC falls off from the NS3C barrel (Fig. 1B). Importantly, the open structures have been observed mostly for linked construct that includes a flexible G4SG4 linker (the disordered residues of NS2B + G4SG4 + the disordered residues of NS3) for the purpose of facilitating protein expression and crystallization (Fig. 1C) ^32,33,39–41^. We use the term “linked” to specifically refer to the G4SG4-linked construct hereafter unless otherwise stated.

The conformational preference of the NS2B/NS3 protease complex can vary significantly among different constructs (“linked” vs “unlinked”) and experimental conditions. Crystal structures of the linked NS2B/NS3 protease predominantly adopt an open state in the absence of inhibitor and a closed state upon ligand binding ^44–48^, whereas solution NMR studies of unlinked and linked constructs of WNV and DENV proteases have shown varying results ^42,47^. One NMR study found that the closed state is the predominant state of the linked WNV protease irrespective of the presence of inhibitor ^47^. However, this NMR study with linked protease has several signal overlaps and line broadening which may lead to biased extrapolation of resonance assignments towards the ligand bound closed state. Another NMR study with a higher resolution showed that the unlinked DENV protease exists predominantly in the close conformation ^48^. This study also argues that unlinked construct is more realistic as the active form of the protease for biochemical study and inhibitor development. In terms of the linker effect, though it can facilitate protein expression and crystallization, biochemical measurements have indicated that linked proteases from DENV and ZIKA exhibit substantially lower catalytic activities under the same physiological conditions compared to the unlinked construct ^49^. Caution has been advised regarding the use of the linked construct of the protease, especially for inhibitor screening ^49^. The underlying mechanism of how the linker affects the structure and dynamics of the protease requires further investigation.

The active site of the NS2B/NS3 protease is formed by the highly charged H51-D75-S135 catalytic triad (residue numbering from PDB: 5IDK ^42^), which normally recognizes substrates with dibasic residues at the P1 and P2 sites and a small residue at the P1’ site ^37^. Substrate-based active-site binding inhibitors would also be highly negatively charged by design, and this leads major challenges in low oral bioavailability ^49^. An attractive alternative could be to target transient folding intermediate states of the protease and/or disrupt the formation of the functional enzymes using allosteric inhibitors ^50,51^. For example, several potent broad-spectrum NS2B/NS3 inhibitors have been discovered by screening a set of 2816 approved and investigational drugs that directly target the NS2B-NS3 interaction interface ^52^. These compounds likely bind to an intermediate, partially folded NS3 conformational state and prevent the binding of NS2B to form the active enzyme complex ^52^. Current searching for allosteric inhibitors has only produced mostly sub-micromolar to micromolar inhibitors ^53–57^. Furthermore, understanding of possible targeting conformational states, allosteric sites as well as molecular mechanisms of allosteric inhibition is limited.

At present, a comprehensive characterization of protease folding pathways is still lacking. Computational simulations have proven a powerful approach for studying protein folding mechanisms and pathway ^58–61^, especially using extremely efficient topology-based models ^60,62–66^. Topology-based models are also known as Gō or Gō-like models. They are based on the consistency principle originally proposed by Nobuhiro Gō ^67^, which posits that all interactions as observed in the native structure are mutually consistent and each independently supports the formation of the native fold ^68,69^. The idea is closely aligned with the “principle of minimal frustration”, arguing that evolutionary selection has shaped protein energy landscapes to be smooth and funnel-like with stabilization dominated by native contacts ^59,60,70^. As such, the free energy landscape of a folded natural protein can be well approximated by considering only the contributions of native contacts, assuming that formation of non-native contacts during folding is minimally stabilizing (i.e., minimal frustration). Importantly, contributions of non-specific interactions can be included in topology-based models to account for the presence of various physical forces on protein folding ^66,71–73^. The success of topology-based modeling is well documented ^60,73–79^. Despite major advances in modern atomistic protein simulations ^80^, topology-based modeling remains a highly valuable tool in modern protein folding study ^81–83^.

In this work, we employed the sequence-flavored Gō model to investigate the coupled binding and folding pathways of the WNV NS2B/NS3 protease complex. The sequence-flavored Gō model is Cα-based and includes sequence-based backbone torsion terms and statistical potentials for native contacts formed by different residue pairs ^71,84^. The sequence flavor allows the model to predict subtle differences in folding mechanisms that arise from sequence differences in topologically analogous proteins ^71^. Calibrated using atomistic simulations, extensive molecular dynamics (MD) simulations using the resulting Gō-like models can provide a holistic view of the folding pathways of both G4SG4-linked and unlinked WNV NS2B/NS3 proteases, revealing detailed conformational intermediate states. Our analysis also explains how the G4SG4 linker destabilizes the NS2B C-terminal hairpin and perturbs the folding pathways, which is consistent with the reduced protease activities measured in previous experimental studies ^85^. Our work represents the first analysis of the folding pathway of orthoflavivirus proteases and provides insights of several critical intermediate states along the folding process, which can potentially help progress characterization of potential allosteric sites and the future design of allosteric antiviral drugs.

## Results

### Calibration of topology-based Gō models for the WNV NS2B/NS3 protease

All-atom structure of the WNV protease in the close state (PDB: 5IDK ^42^) was used to derive the native contact pairs and generate a Cα-based sequence-flavored Gō model using the MMTSB Gō Builder ^71,86^ (see Methods). Benchmark simulations at 300 K reveal that the original Gō model was too weak, unable to maintain the structures of NS2B, NS3 or the complex (Figure S1). This is also reflected in the very large root mean square fluctuation (RMSF) profile (Fig. 2A). We calibrated the model by uniformly scaling up the Gō potential by a factor of 1.14 to reproduce the RMSF profile derived from 500 ns all-atom simulations of the unlinked NS2B/NS3 (Fig. 2A blue vs black traces). With the scaling, the model was able to maintain a stable folded structure during the control simulations at 300 K (Fig. S1). The contact map of the final calibrated Gō model shows the beta-structure patterns of the protease complex and the range of native-contact potential strengths and distances (Fig. 2B, 2C), which was used for all further simulations in this work.

**Figure 2.**
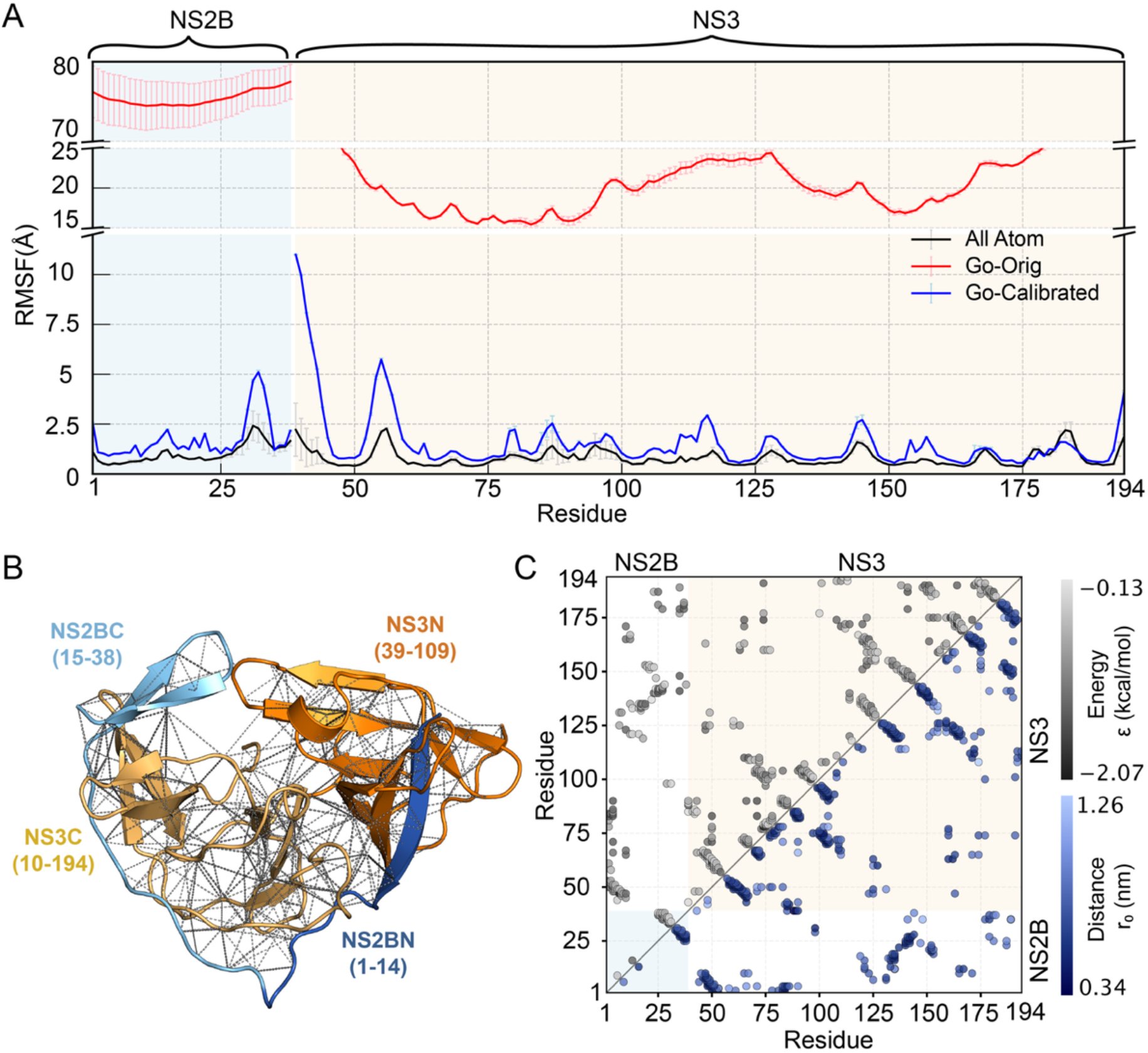
Gō model calibration for the NS2B/NS3 complex. **A)** RMSF profiles of the unlinked construct simulated using the original and final calibrated Gō models, in comparison to the one derived from all-atom explicit solvent simulation using CHARMM36m. The mean and standard error are calculated from 4 independent simulations. **B)** All 308 native contacts (dashed lines) of the NS2B/NS3 protease complex, with the color gradient of gray representing the strengths of the Gō potentials (see color bar in panel C). **C)** Contact map showing the strengths (upper triangle) and Cα distances (lower triangle) of the 308 native contact pairs.

### Gō modeling recapitulates the destabilizing effects of the G4SG4 linker

An intriguing observation from structural studies is that the open state with NS2BC hairpin unbound from NS3 (Fig. 1B) has been mostly observed with the G4SG4 linker construct ^39,43,43,45,87,88^, while solution NMR studies showed that the protease remains in the closed or nearly closed state for unlinked WNV NS2B/NS3 protease even without inhibitor ^48,89^. Other studies further demonstrated that the linker can destabilize the catalytic active complex conformation and reduce the protease activity ^85,90^. Indeed, simulations of the G4SG4 linked construct using the calibrated Gō model shows that the NS2BC hairpin is unstable and tends to unbind from the complex (Fig. S2), while the rest of the complex (NS2BN and NS3) displays a similar level of RMSF values compared to that from the unlinked construct or the all-atom simulations (Figure 3). Note that the disordered G4SG4 linker is only 30 residues long, including the 8-residue and 13-residue disordered tails of NS2B (residues 39-46) and NS3 (residues 56-68) (see Methods for residue numbering scheme). Treating the entire disordered segment as a Gaussian chain ^91^, the mean end-to-end distance can be estimated as 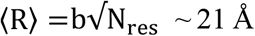, with the residue number N_res_ = 30 and the Cα-Cα distance b ∼ 3.8 Å. Therefore, the linker needs to adopt a relative extended conformation to bridge the ∼50 Å separation between the NS2B C-terminus and NS3 N-terminus in the structured region (see in Fig. 1C). The entropy cost associated with stretching the linker results in the destabilization of the NS2BC hairpin. To corroborate this hypothesis, we constructed another linked NS2B/NS3 construct with 16 G4SG4 repeats, termed as (G4SG4)_16_, such that the mean end-to-end distance of the new 169 residues linker, including 21 disordered residues from NS2B/NS3 termini, 144 residues from (G4SG4)_16_, and 4 additional G, would match the ∼50 Å separation between the two termini in the structured region. The results show that the long linker construct with the same calibrated Gō model has similar RMSF profile compared to the unlinked construct, with no destabilization of the NS2BC hairpin (Fig. 3, Fig. S2). The ability to capture the linker effects on the stability of the complex further supports that the calibrated Gō model properly balances native interactions and conformational flexibility and is suitable for studying the mechanism and pathways of the coupled binding and folding of the NS2B/NS3 protease.

**Figure 3.**
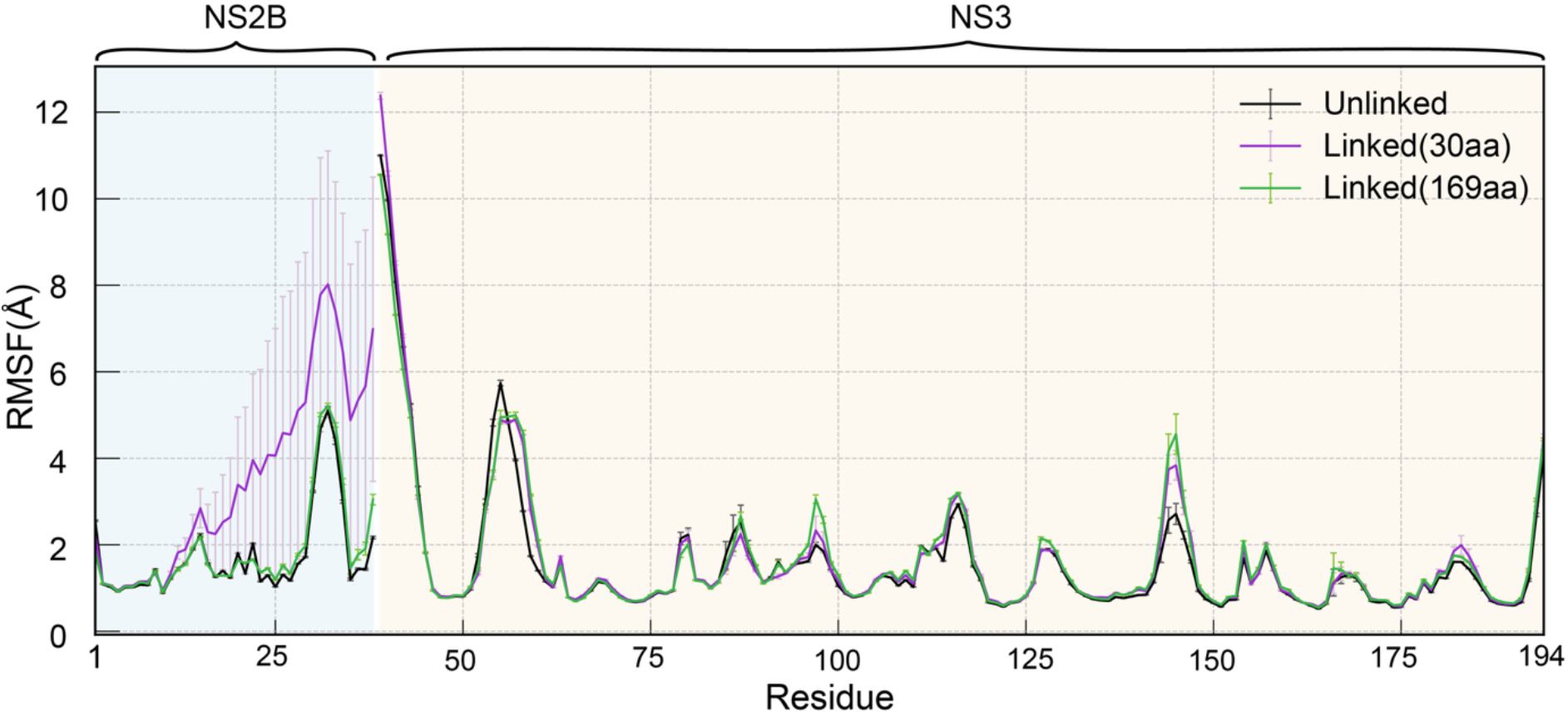
RMSF profiles of different NS2B/NS3 constructs simulated using the calibrated Gō model. The unlinked, linked constructs (G4SG4 and (G4SG4)_16_) constructs are colored black, purple and green, respectively. Each RMSF was averaged over 4 replicas shown by the mean standard error bars.

### Parallel folding pathways and multiple intermediate states in NS2B/NS3 complex formation

Long MD simulations have been previously performed near the melting temperature (T_m_) to sample multiple reversible binding and folding transitions to analyze the mechanism and pathways of intrinsically disordered proteins (such as NS2B here) ^66,92–94^. Exploratory simulations suggest that folding of double barrels in the protease complex structure is highly cooperative and a precise determination of T_m_ where reversible binding/folding transitions could be directly sample proved not feasible. We failed to observe reversible transitions within 10 µs for all temperatures tested including those shown in Fig. S3. As such, we focused on refolding simulations starting from an ensemble of 16 unfolded structures derived from high temperature simulations (Fig. S4, Fig. S5, see Methods). Four independent 10-µs simulations at 300 K were performed for each initial configuration using the calibrated Gō model. From the 64 refolding trajectories, we observed the folding transitions in 51 runs for the unlinked construct and in 47 runs for the linked construct (Fig. S6), as shown by the fraction of the native contacts between the NS2B/NS3 interface. All folding transition segments were collected for subsequent mechanistic analysis (see Methods).

The conformational space network (CSN) analysis was employed to visualize the conformational states and transitions during the formation of the native complex ^66,95–97^. Here, the network was constructed using a graph representation where each node represents a conformational microstate, and edges define connectivity reflecting transitions between microstates (nodes) observed in simulations. Since our study uses topology-based models, a natural choice for the order parameters to define conformational microstates is native contact fractions involving different segments of the protease. Specifically, we defined four natural segments of the protease: NS2BN (residues 1-14), NS2BC (residues 15-38), NS3N (residues 39-109), and NS3C (residues 110-194), and calculated 9 intra- and inter-segment native contact fractions, namely, 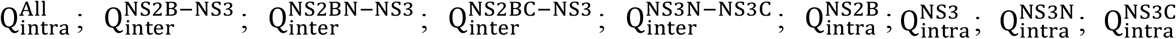 (also see Table 1). Each native fraction ranges from 0 to 1 and is separated into five equidistant bins for indexing each snapshot sampled in folding transition segments (see Methods).

**Table1.** Definition of NS2B/NS3 segments and numbers of various intra- and inter-segment native contacts.

| Type of interactions | Segment(s) | Residues | # of native contacts |
| --- | --- | --- | --- |
| Intra | All | 1-194 | 308 |
| Intra | NS2B | 1-38 | 12 |
| Intra | NS3 | 39-194 | 210 |
| Intra | NS3N | 39-109 | 80 |
| Intra | NS3C | 110-194 | 103 |
| Inter | NS2B/NS3 | 1-38/39-194 | 86 |
| Inter | NS2BN/NS3 | 1-14/39-194 | 44 |
| Inter | NS2BC/NS3 | 15-38/39-194 | 42 |
| Inter | NS3N/NS3C | 39-109/110-194 | 27 |

The resulting CSN derived from the ensemble of folding transitions for the unlinked protease shows six major intermediate state clusters (I1-I6) in addition to the unfolded (U) and folded (F) states (Fig. 4A and Fig. S7). As shown in the CSN, coupled binding and folding of the unlinked protease can proceed in two possible branches: the I1 state, where the NS3C barrel folds whereas no other parts of the complex have yet folded, and the I2 state, where the NS3N barrel is largely folded with NS2BN bound whereas other parts are unfolded. Both states I1 and I2 can proceed to the I4 state where both NS3N and NS3C barrels are formed independently, and NS2BN is bound to NS3N. However, the interface of the two NS3 barrels has not yet formed. The I1 state can also evolve to a separate state I3 beside I4, where NS2BC folds into a hairpin and binds to the already-formed NS3C barrel. Both the I3 and I4 states could evolve to state I5, which has well-folded NS3N and NS3C barrels bound to NS2BN and NS2BC respectively while the barrel interface is not formed. State I4 can also proceed to state I6, which has the whole NS3 domain folded with NS2BN bound but not NS2BC. The network overall shows multiple parallel folding pathways where the protease can go through several intermediate states before reaching the fully bound and folded state (F). The CSN derived from folding trajectories of the linked protease is generally consistent with that of the unlinked protease, except that intermediate state I3 is absent (Fig. 4B and Fig. S8). This will be further discussed later.

**Figure 4.**
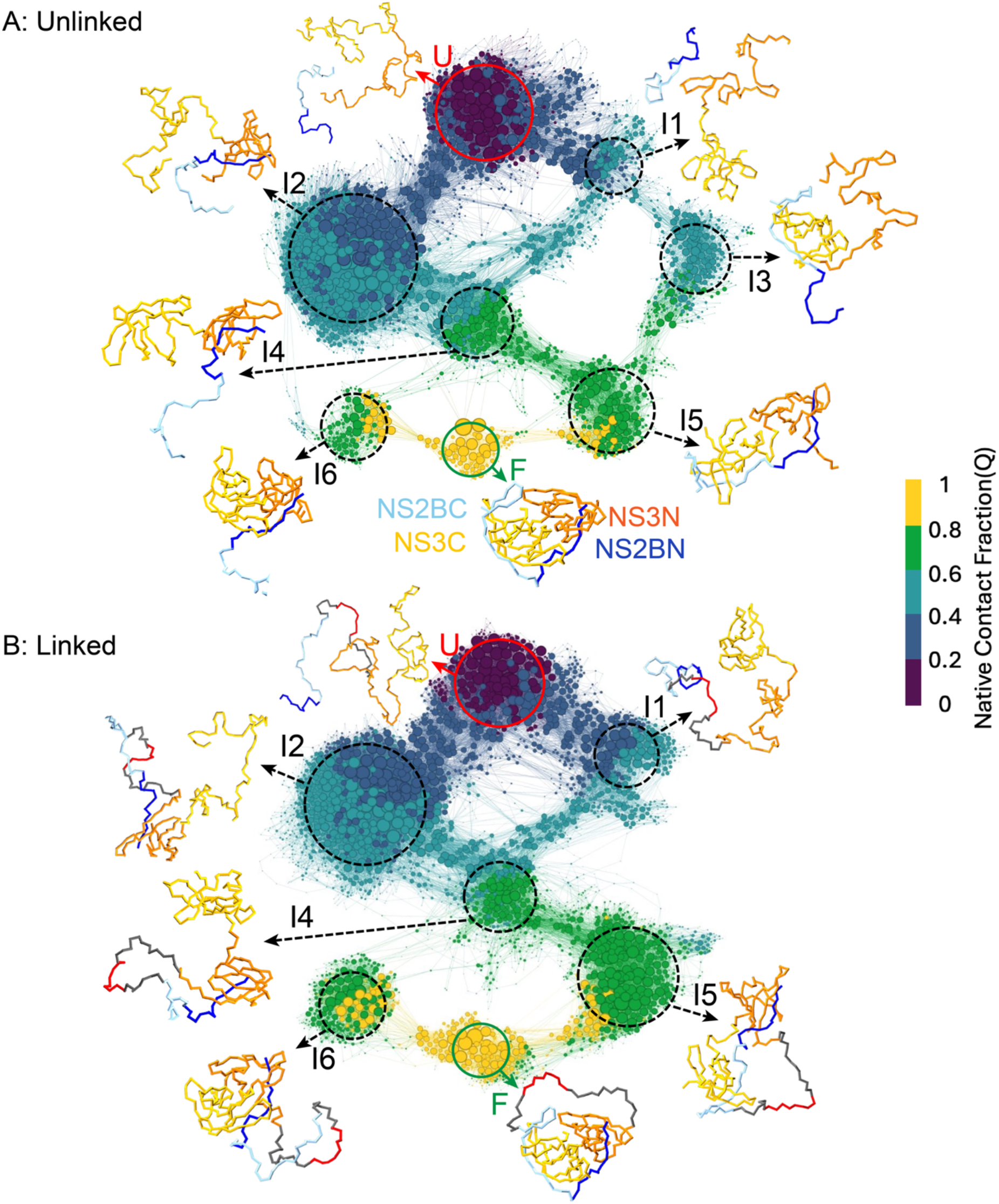
CSN of the folding of NS2B/NS3 complexes. **A)** The unlinked construct; and **B)** the linked construct. The node size reflects the logarithmic scale of the statistical weight of the conformational microstate, and the edge width represents the number of transitions observed. The CSN is colored by 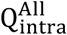, which reflects the overall folding progress. Representative conformations are shown for each state, colored using the same scheme as shown in Fig. 1. Additional representations of these CSNs are given in Figures S5 and S8 with nodes colored using each of all 9 native contact fractions defined in Table 1.

### Retracing and partial folding-unfolding transitions along multiple pathways

Observations of many parallel folding pathways (and intermediate states) can be verified by inspection of individual folding transitions over time. Representative folding trajectories of the unlinked construct are shown in Fig. 5 using 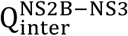 and 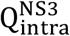 as the two main order parameters to capture the progress toward the formation of the folded complex (see Figs. S9 and S10 for all trajectories of both linked and unlinked constructs). These trajectories illustrate the diversity of folding pathways and how the system may visit various intermediate states during the folding process. Interestingly, some trajectories sample intermediate states along one pathway before retracing and proceeding along another towards the folded complex (e.g., Fig. 5: s62, Fig. S9: s35, s36, s58). Although we did not directly observe the folding-unfolding process of the whole protease, we observed numerous partial folding-unfolding transitions among intermediate states as well as between them and the unfolded and folded states. For example, Fig. 5: s41 shows the folding-unfolding transition between the partially folded I6 state and the folded state (F), which involves the NS2BC binding to NS3 or peeling off. Folding-unfolding transitions were also observed between I2 and I4 states (e.g., Fig. 5: s3, Figure S9: s28, s35), as well as between the unfolded state (U) and I2 state (e.g., Fig. S6: s60). Furthermore, Fig5: s2 shows an example of direct and highly cooperative transition from partially folded I2 state to the fully folded state (F). Although many folding pathways and intermediate states are shared in both the unlinked and linked protease folding, we observed that the I3 state (Fig. S6: s9-11, s58-59), which are traversed during unlinked protease folding, is complete absent in the linked folding trajectories (Fig. S10, Fig. 4B). The presence of the linker makes it highly unfavorable for NS2BC hairpin folding (Fig. 2), thus preventing the formation native NS2BC/NS3C interface characteristics of state I3.

**Figure 5.**
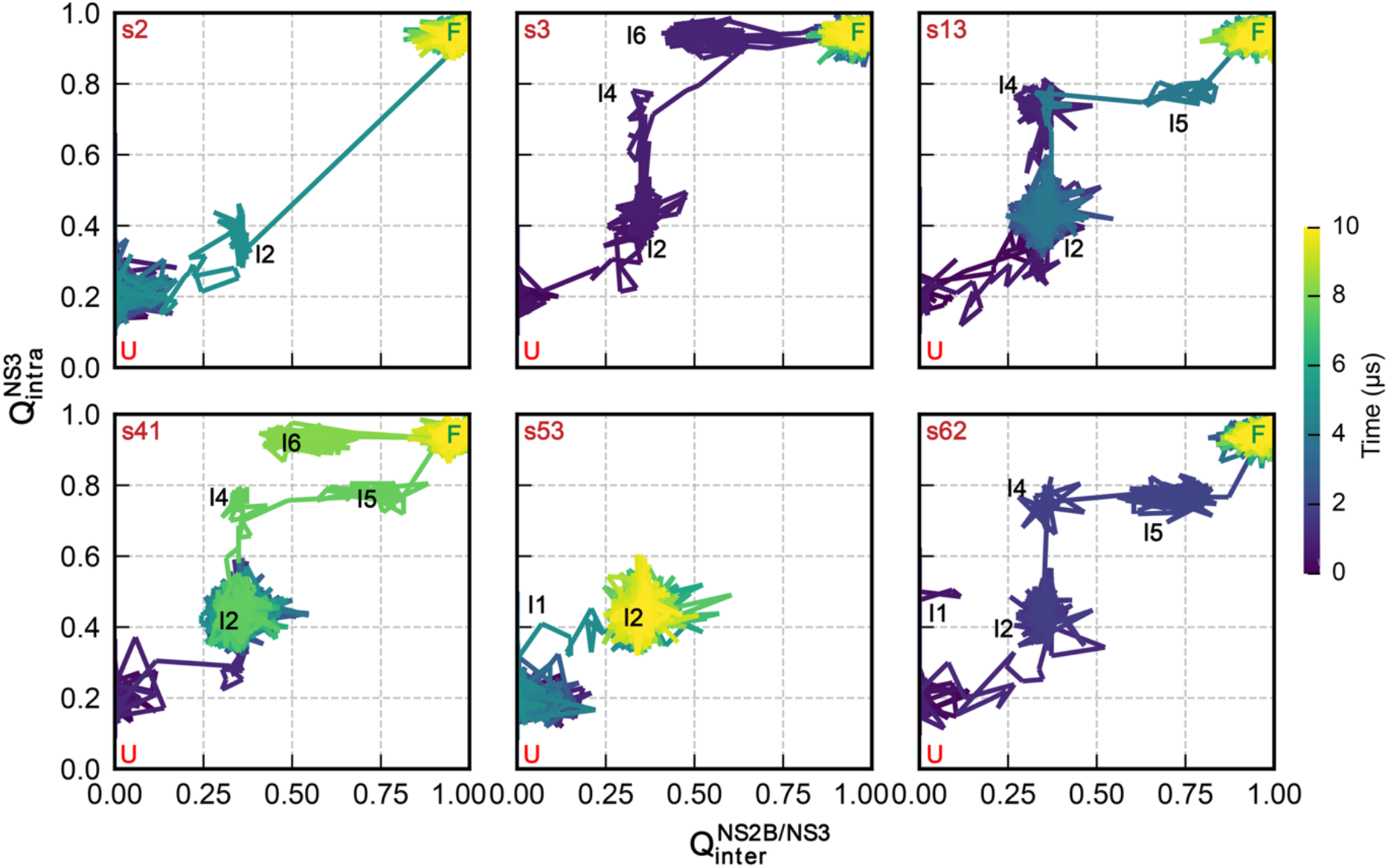
Representative folding transition trajectories of the unlinked construct. The folding trajectories are characterized using 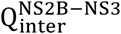 and 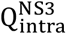 and colored using the simulation time. Intermediate states visited during each trajectory are also marked.

### Pseudo-free energy landscapes of NS2B/NS3 folding: critical importance of NS3N–NS3C interface formation

We further constructed 2D pseudo-free energy landscapes along various combinations of native contact fractions (Table 1) using the folding transition segments collected from 64 independent folding simulations to study protease folding mechanisms (see Methods). Since the folding of protease involves the binding of NS2B and NS3 to form the complex and NS3 is the core domain that encompasses two barrels, we selected 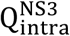 and 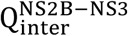 as two of the most informative order parameters. As shown in Figure 6 A&B, intermediate states identified through CSN analysis map well into different local free energy minima, with the only exception of the I3 state, which overlaps partially with the I2 state due to dimensional reduction. Folding of both unlinked and linked constructs is a multi-step and multi-path process that involves both “induced folding” and “conformational selection” mechanisms (Fig. 6A, 6B) ^98–100^. In particular, the folding of the two NS3 barrels exhibits different mechanisms. NS3N barrel folding is strongly coupled to NS2BN binding, following a largely cooperative coupled binding and folding mechanism (Fig. S11A, S12A), whereas NS3C barrel folding can occur independent of NS2B binding, following a conformational selection-like mechanism (Fig. S11B, S12B). Note that despite NS3C itself can fold, the folded state of NS3C alone is less stable unless bound to NS3N to form the NS3 core domain (Fig. S11E, S12E), which is consistent with the observation that both I1 and I4 are less stable compared to the unfolded (U) and I2 states (Fig. 4; Fig. 6A, 6B). This also explains why the bottleneck of NS3N/NS3C interface formation resulted in more populated states in state I2 instead of I4. The changed contour around I2 states shows the missing of I3 states in the linked protease folding pathway (Fig. 6A, 6B; Fig. S11F, S12F).

**Figure 6.**
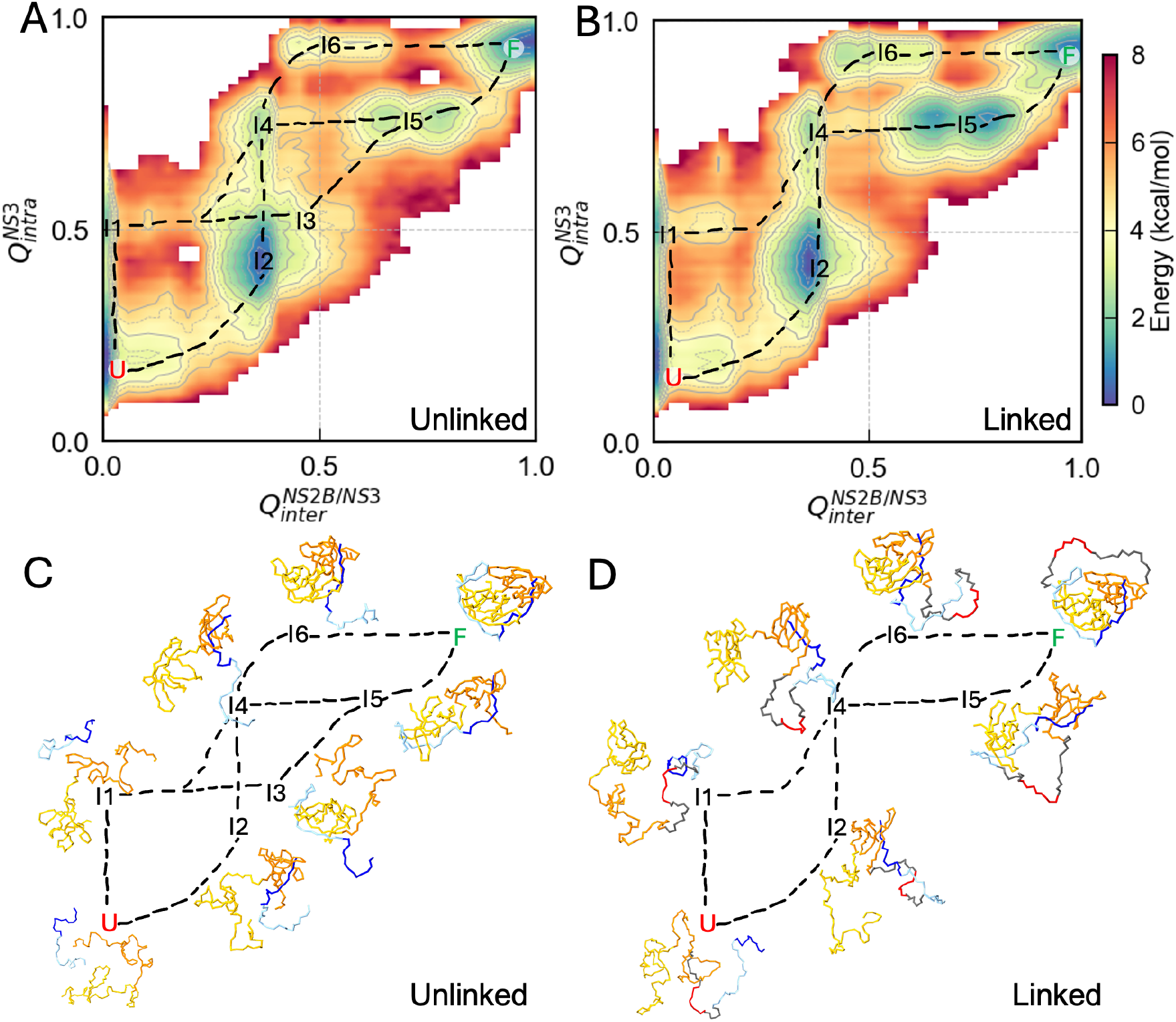
2D pseudo-free energy landscape of the folding along two native contact fraction order parameters: 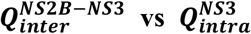. **A)** and **B)** pseudo-free energy landscapes of the unlinked and linked constructs represented via potential mean force calculated based on the native contact fraction probability. **C)** and **D)** Representative conformational states projected on the 2D pseudo-free energy landscape for the unlinked and linked construct. All folding pathways are annotated by connecting the mapped low free energy states using dashed lines.

The dominant bottleneck during protease folding is associated with formation of the interdomain interface between NS3N-NS3C β-barrel domains. From CSN analysis, we observed fewer and thinner edges connecting states I4 to I6 and states I5 to F (folded state), which highlights the folding bottleneck as the formation of the NS3N/NS3C interface in with both NS2BC bound (I5) or unbound (I6) states (Fig. 4). Among the 6 identified intermediate state clusters, the intermediate states I2 and I5 are identified as the most highly populated and highly connected node clusters in the folding network of both the unlinked and linked constructs (Fig. 4; Fig. 6A-B). Although the I1 and I4 states are also essential in the folding path, they are less populated and more transient possibly due to the folded NS3C barrel alone is unstable as previously mentioned and unfolds back to unfolded state (U) or I2 state clusters. This is supported by the dense edges between I2 and I4 state clusters and the dense edges between U and I1 state clusters (Fig. 4). The importance of I2 and I5 states is supported by the 2-D trajectory evolution analysis which shows that most successful folding trajectories traversed I2 state (42/51 for unlinked, 43/47 for linked, Fig. S13-14) and I5 state (37/51 for unlinked, 34/47 for linked, Fig. S15-16). Interestingly, these two state clusters are also intermediate states where the unsuccessful folding trajectories were trapped, where NS3 inter-barrel interface failed to form to allow folding to proceed (Fig. S17-18). Although these simulations do not sample fully reversible folding–unfolding transitions and therefore do not permit quantitative estimation of free-energy barriers, the pronounced accumulation of folding flux and prolonged residence at states I2 and I5 identifies the NS3N/NS3C interface formation as dominant bottleneck along the forward folding pathways. The result suggests these two states may be the main metastable intermediate states that could be targeted for allosteric inhibitors of the protease.

## Discussion

The orthoflavivirus protease, NS2B/NS3, has been considered as a promising antiviral drug target to treat life-threatening diseases related to orthoflavivirus infection. Yet, lack of molecular understanding for its folding process constraints drug development, especially for allosteric inhibitors, where information of folding intermediate states would be greatly beneficial. Our calibrated Gō model for the protease can not only reproduce structural dynamics seen in atomistic simulations (Fig. 2A) but also recapitulate the destabilization effect from the G4SG4 linker used in many previous experiments (Fig. 3). This finding resolved the previously puzzling role of the G4SG4 linker in the protease structure and function reported in different studies ^47–49,93,94^. Large-scale re-folding simulations initiated from diverse sets of unfolded configurations successfully reveal a holistic picture of the protease folding landscape, major intermediate states, as well as detailed coupled binding and folding mechanisms.

The sampled folding landscapes for the unlinked and linked construct share highly similar features. The protease can initiate its folding independently from either NS3N or NS3C barrels folding, though the NS3N folding is more favorable due to the coupled binding with NS2BN. This renders the I2 state (with only the NS3N folded and NS2BN bound) to be the dominant intermediate state. Furthermore, formation of the NS3N-NS3C barrel interface has emerged as the major folding bottleneck. The N-terminal barrel formed by the NS2BN/NS3N that exists in I2, I4, I5 states could be considered as an essential conformation of the folding intermediate states that can be potentially targeted for novel allosteric protease inhibitors. Interestingly, the G4SG4 linker has important implications on the folding landscape besides its destabilization effect on NS2BC hairpin in the folded state. The linker largely eliminate binding and folding of NS2BC to NS3C (state I3) in the linked construct prior to the folding of the rest of the protease (Figures 4 and 6). Nonetheless, the linker does not seem to affect the location of the main folding bottleneck.

The NS3 core domain alone can fold under the guidance of the native-contact potential which is not reflecting the real case in cell (Fig. S19). This observation suggests that the calibrated Go-like potential used in this study may be stronger than the original native-contact potential. This reflects an inherent trade-off between the computational timescale required to observe folding transitions and the physical accuracy of the model. A more finely tuned potential that promotes folding of the NS2B/NS3 complex—but not the isolated NS3 domain— would likely require substantially longer simulation times to capture folding events. Therefore, we adopted this calibrated Gō potential as a practical compromise, acknowledging that it may slightly over-stabilize the folded state. Despite the over-stabilization, the model shows that NS3 core in the NS3-alone simulation has several more flexible loops compared the NS3 in the NS2B/NS3 complex dynamic simulation which have previously been found to be related to the allosteric regulation of the protease catalytic function (Fig. S20) ^38^.

Despite the encouraging folding pathways derived from the simple topology-based model, several inherent limitations need to be noticed for such models. First, the details of the unbound states cannot be fully represented by the Cα based Gō model due to the nature of structure-based model. Second, non-native contact does not exist in the final folded state structure, however it may involve in the folding transition by providing additional energetically accessible pathways that can lead to the folded native state ^103–105^. Third, this structure-based model focused on the protease complex alone and we did not consider the potential role of chaperons playing in the folding process of flavivirus protease folding ^106–108^. Finally, the tradeoff between simulating the folding transition time and the accuracy of Gō potential strength need to be considered when interpreting our results.

In conclusion, our study revealed the mechanisms of the orthoflavivirus WNV NS2B/NS3 protease folding processes, the effect of linker on the protease dynamics and folding process. We identified the several intermediate states and pathways as well as the major folding bottleneck of the protease folding process which can provide insights for the drug design.

## Methods

### Gō model generation and calibration

All-atom structure of NS2B (resid: 51-88) segment and the NS3 (resid: 1014-1169) segment were extracted from PBD: 5IDK ^42^ and renumbered as NS2B: 1-38 and NS3: 39-194. Hydrogens were added to the cleaned structure with an in-house written CHARMM script, which was uploaded to the MMTSB Gō Builder to generate the sequence-flavored Gō model ^63,71,109,110^. The initial Gō model was further calibrated by uniformly scaling all native contact pair potentials to reproduce the Cα RMSF profile from atomistic simulations (see below). A scaling factor of 1.2 was chosen, which yields a T_m_ of around 340 K. This model also allows sampling of folding transitions at 300 K within a feasible timespan (∼10 µs in this work). To model linked constructs, two linkers of different lengths were added to the all-atom structure using ColabFold server ^111^. The first 30-aa linker includes G4SG4 used in experimental studies as well as the disordered segments (residues 39-46 and 55-68) from the protease connected to NS2BC or NS3N; the second 169-aa linker includes (G4SG4)_16_ repeats and the disordered tails. In the linked Gō models, the linker is considered fully disordered and is not involved in any native contacts. In other words, the same set of NBFIX native contact potentials are used in the unlinked and linked Gō models.

### Gō model simulations

All Gō model simulations were performed using CHARMM ^112^. The Gō models were energy minimized with steepest descent and Adopted Basis Newton-Raphson method and then equilibrated with weak harmonic positional restraints imposed. The production runs were performed using Langevin dynamics with a timestep of 0.01 ps and collision frequency of 0.1 ps^-1^. For model calibration, 4 independent simulations were performed for 5 μs at 300 K and 350 K for each system (unlinked, linked with G4SG4, and linked with (G4SG4)_16_), starting from the folded configurations. For refolding simulations, we randomly selected 16 unfolded snapshots from the 350 K trajectories (Figure S19) to initiate 4 independent runs each at 300 K. The refolding simulations lasted 10 μs each, to allow the observation of sufficient refolding events. In total, 64 replicas were run for both unlinked and linked (30-aa linker) constructs.

### All-atom MD simulations

All-atom MD simulations were performed using GROMACS 2019.4 ^113^ with the CHARMM36m force field ^114^. The NS2B/NS3 complex was solvated using TIP3P water ^115^ and neutralized using 0.15 M NaCl using the CHARMM-GUI web server ^116,112,117^. The final cubic system has a dimension of 80 Å with ∼48,000 atoms. Energy minimization was carried out using the steepest descent algorithm for up to 5000 steps, with harmonic positional restraints imposed on protein heavy atoms. The force constants were 400 and 40 kJ·mol^−1^·nm^−2^ on backbone and side chain heavy atoms, respectively. Long-range electrostatic interactions were computed using the particle mesh Ewald (PME) method ^118,119^ with a real-space cutoff of 1.2 nm, while van der Waals interactions were calculated using a force-switch scheme between 1.0 and 1.2 nm. The Verlet cutoff scheme was used for neighbor list construction ^120^. All bonds involving hydrogen atoms were constrained using the LINCS algorithm ^121^. Equilibration was performed under NVT conditions (constant particle number, constant volume and constant temperature) for 125 ps, with the temperature maintained at 303.15 K via the Nosé–Hoover thermostat ^122,123^. The production simulation was performed in the NPT ensemble (constant particle number, constant pressure and constant temperature) for 500 ns using a 2-fs time step, with pressure maintained at 1 bar using the Parrinello–Rahman barostat ^124,125^.

### Structure, native contact and CSN analysis

All structural and contact analysis was performed using the MDanalysis package ^126^. For RMSD and RMSF calculations, trajectories were first aligned to the NS3 core domain using Cα atoms for both all atom and Gō model simulations. Residue numbers of the linked constructs NS3 domain are renumbered to match the unlinked residue numbers (39-194) for the convenience of comparison. Native contacts are considered formed whenever the Cα distance between two residues are no more than 2 Å larger than the native distance. The list of native contacts was provided in the Qlist file generated from the MMTSB Gō builder. The native contacts were further partitioned into various inter- and intra-segment sets as summarized in Table 1. These native contact fractions were used as order parameters for the later CSN and free energy analysis.

To construct the pseudo free energy surfaces and CSNs, we first isolated folding transitions from the set of 64 refolding trajectories, by including from beginning of simulation to no more than 1 μs after the completion of folding. The pseudo free energy surfaces were then directly derived from the 2D histograms of various combination of inter- and intra-segment native contact fractions 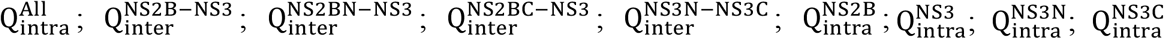; see Table 1), with Gaussian smoothing applied. To construct the CSN, each native contact fraction dimension was binned equally into 5 intervals spanning values from 0 to 1, resulting a total of 5^9^ = 1,953,125 possible conformational microstates (nodes). Each frame from the folding transition trajectories was indexed and assigned to one of the nodes, and an edge was added whenever the system jumped between nodes during the simulations. Visualization of the CNSs was performed via the python package NetworkX ^96,97^. The nodes and edges in the final graphs were stress minimized using the Fruchterman-Reingold for overall layout display of major node clusters and ForceAtlas2 for fine tuning the layout of small node clusters for optimal visualization ^127–129^.

## Acknowledgements

This work is supported by NIH through grants R01 AI156187 (to M. Chen and J. Chen).

## Author contributions

J. Chen and M. Chen conceptualized the project. K. Dong conducted computational modeling. simulation, and analysis. K. Dong, J. Huang, and J. Chen interpreted data, drafted and revised the manuscript. M. Chen reviewed and revised the manuscript.

## Additional information

**Extended data** is available for this paper at https://github.com/kairongdong/p2_ns2bns3_folding_path_GO.

## Supplementary information

The online version contains supplementary material available at https://doi.org/xxxx

