## Supplemental tables and figures for "Coupled Binding and Folding of NS2B/NS3 Protease and Linker Effects Revealed by Topology-based Modeling"

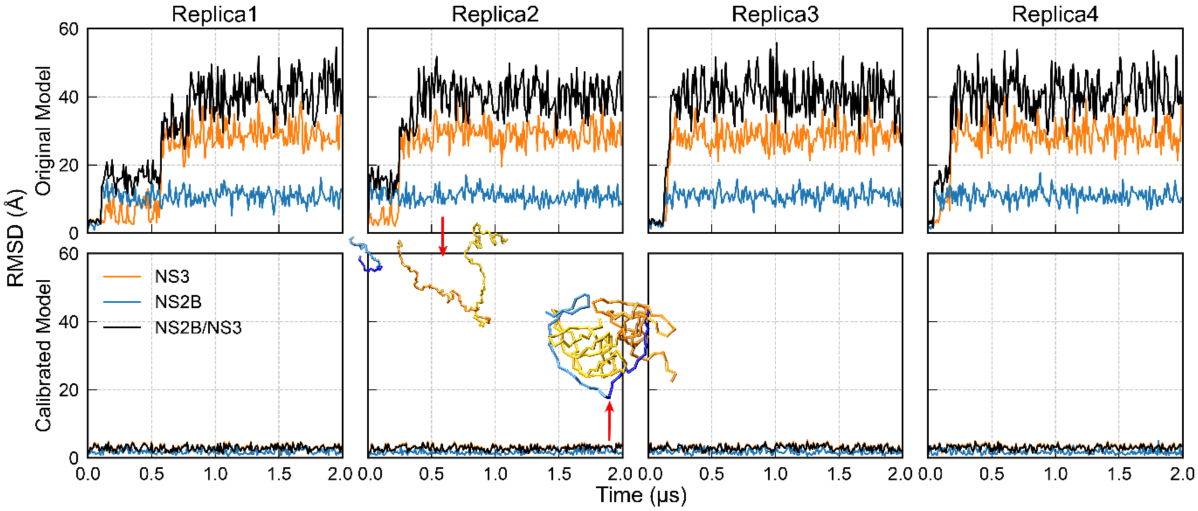


**Supplementary Figure 1. Stability of Gō models for simulating the unlinked NS2B/NS3 protease complex**. The top and bottom rows show the RMSDs of four intendent simulations at 300 K using the original and calibrated NS2B/NS3 Gō models. Representative snapshots are shown in trace representations with NS2B in blue and NS3 in orange.

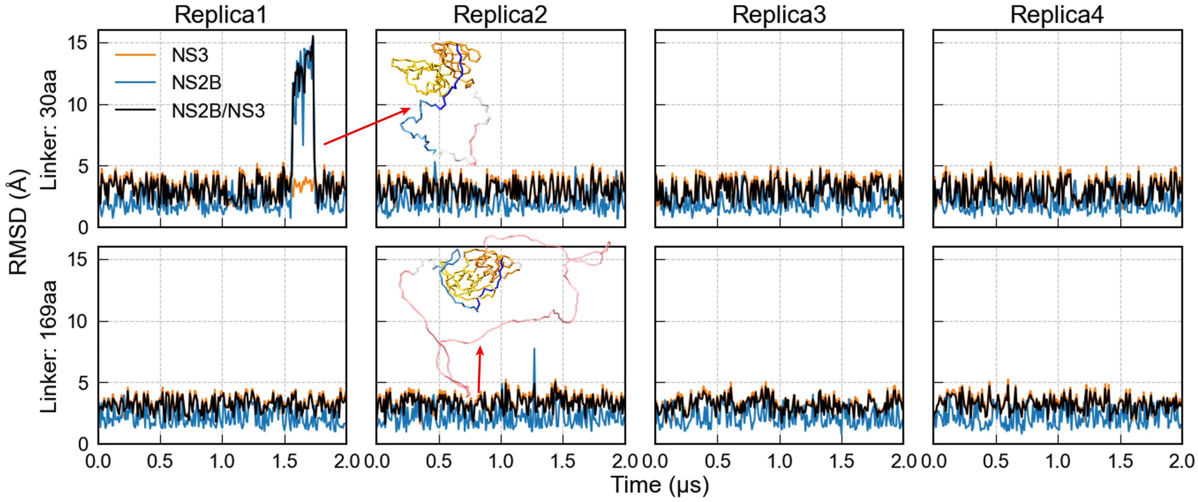


**Supplementary Figure 2. Stability of two linked constructs of NS2B/NS3 protease simulated using the calibrated Gō model**. The top and bottom rows are for constructs with the $G_4SG_4$ and $(G_4SG_4)_{16}$ linkers, respectively. All simulations were performed at 300 K. Representative snapshots are shown with the linker shown in pink.

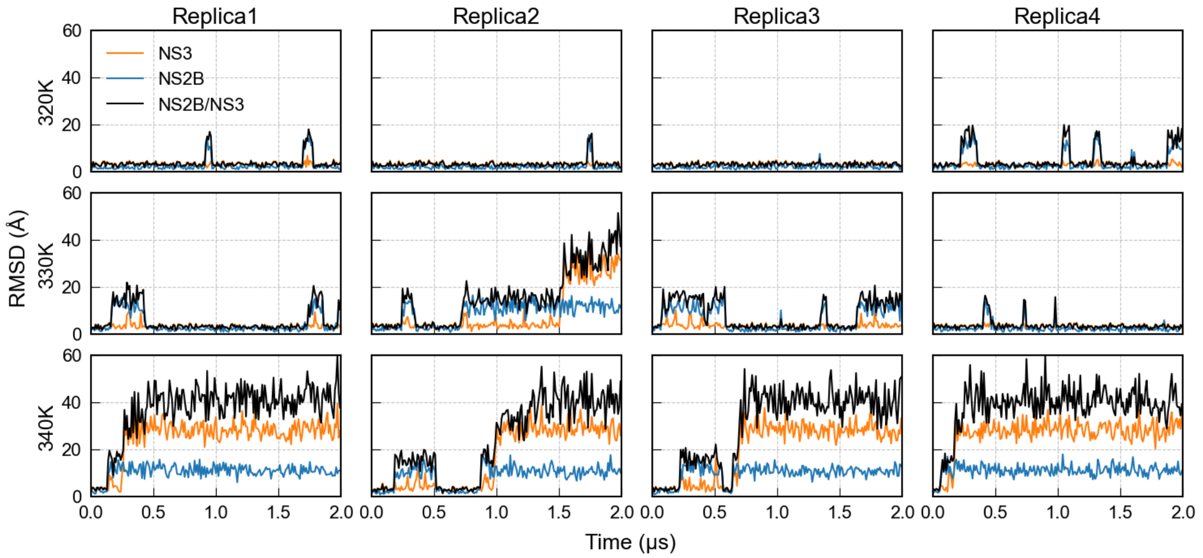


**Supplementary Figure 3. Simulations of unlinked NS2B/NS3 at 320 K, 330 K and 340 K.** The RMSD trajectories of NS2B, NS3, and the complex were calculated separately using the initial folded state and shown in orange, blue and black, respectively.

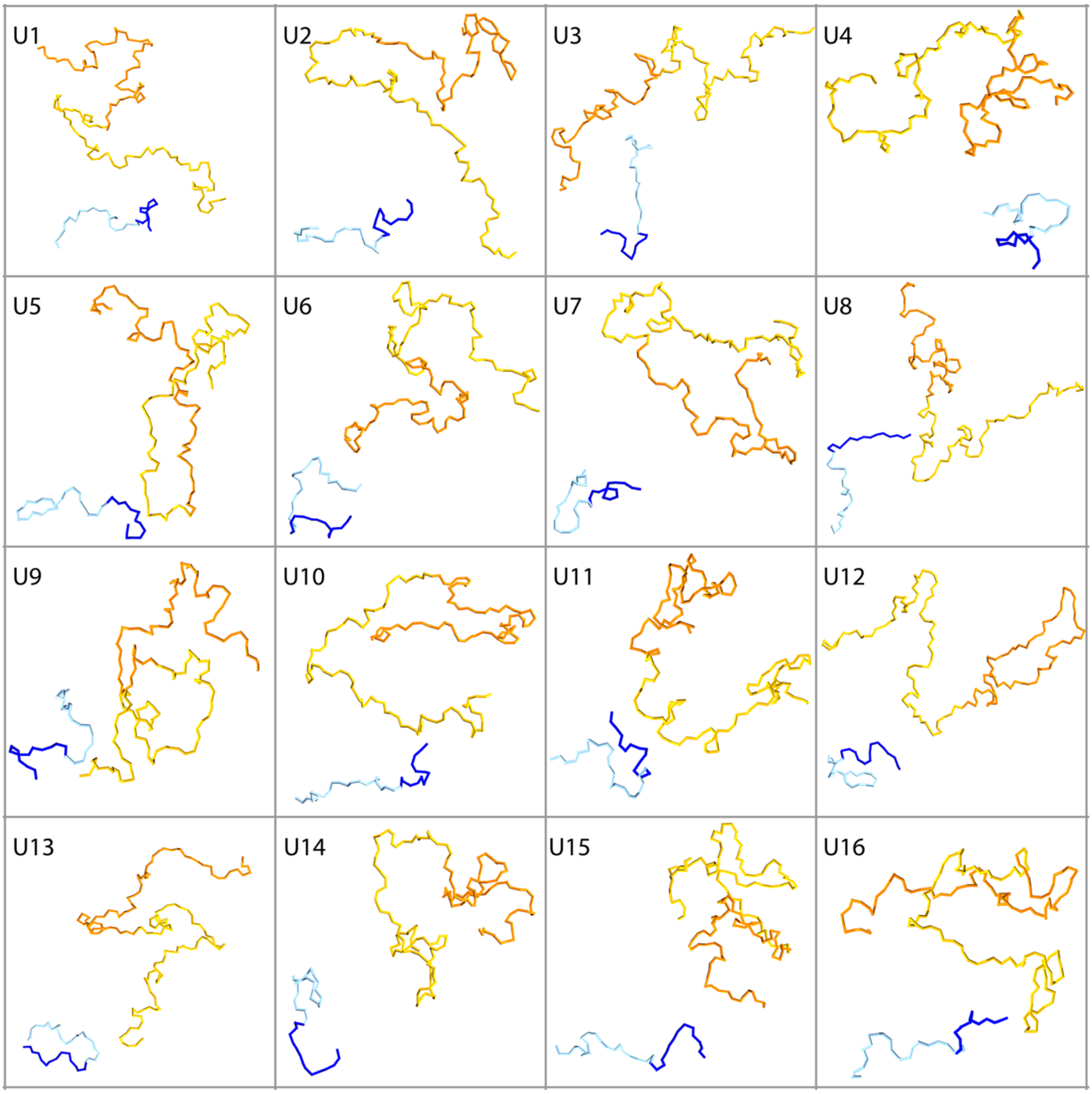


**Supplementary Figure 4. The 16 selected unfolded conformations used as the initial conformations for folding study of the protease (unlinked).** NS2BN (resid 1-14) is colored with dark blue; NS2BC (resid 15-38) is colored with light blue; NS3N (resid 39-109) is colored with orange; NS3C (resid 110-194) is colored with yellow.

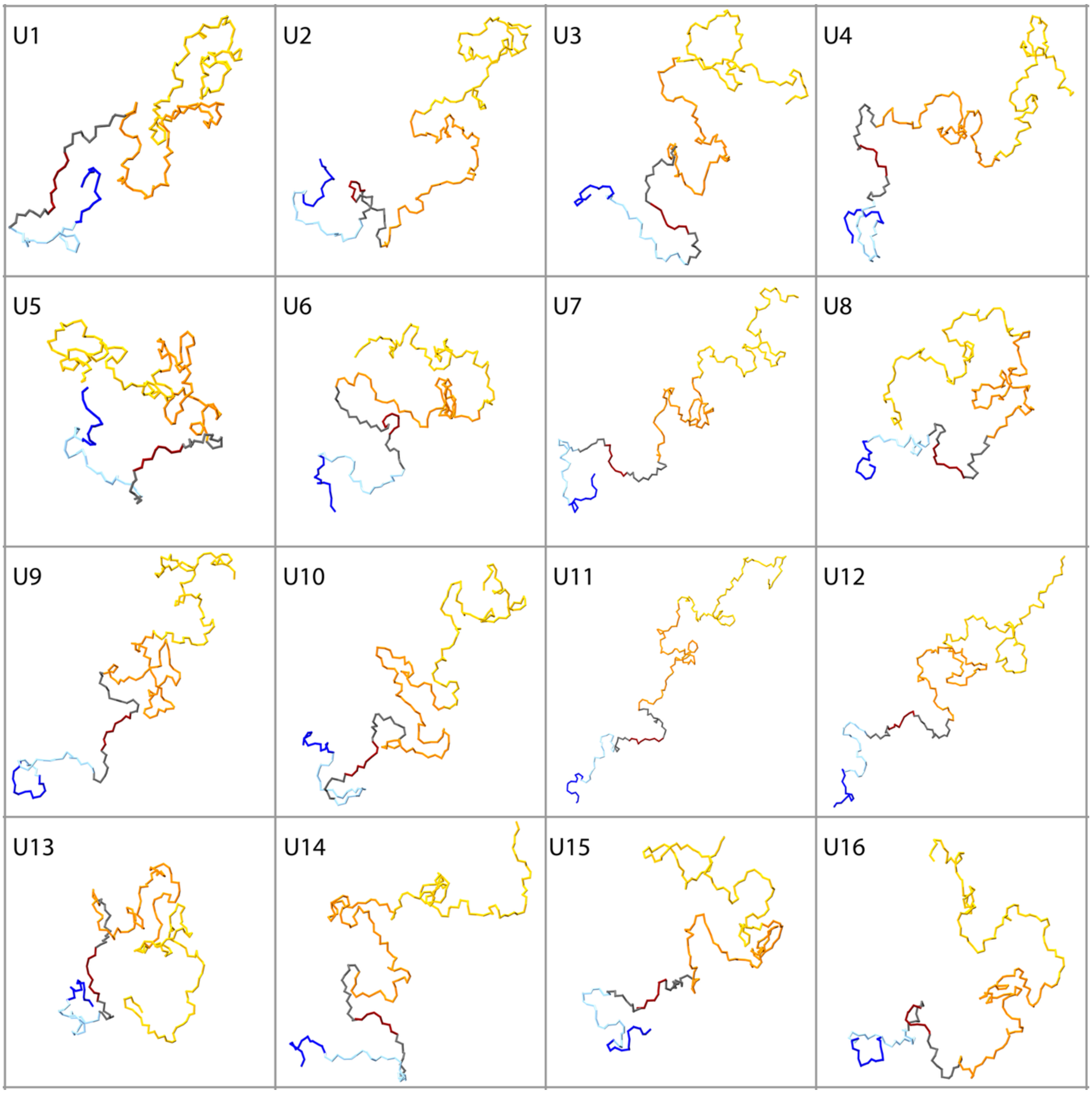


**Supplementary Figure 5. The 16 selected unfolded conformations used as the initial conformations for folding study of the protease (linked with G4SG4).** NS2BN (resid 1-14) is colored with dark blue; NS2BC (resid 15-38) is colored with light blue; NS3N (resid 69-139) is colored with orange; NS3C (resid 140-224) is colored with yellow; G4SG4 linker (resid 47-55) is colored with red; the disordered residues (resid 39-46 and resid 56-68) connecting with NS2BC and NS3N are colored gray.

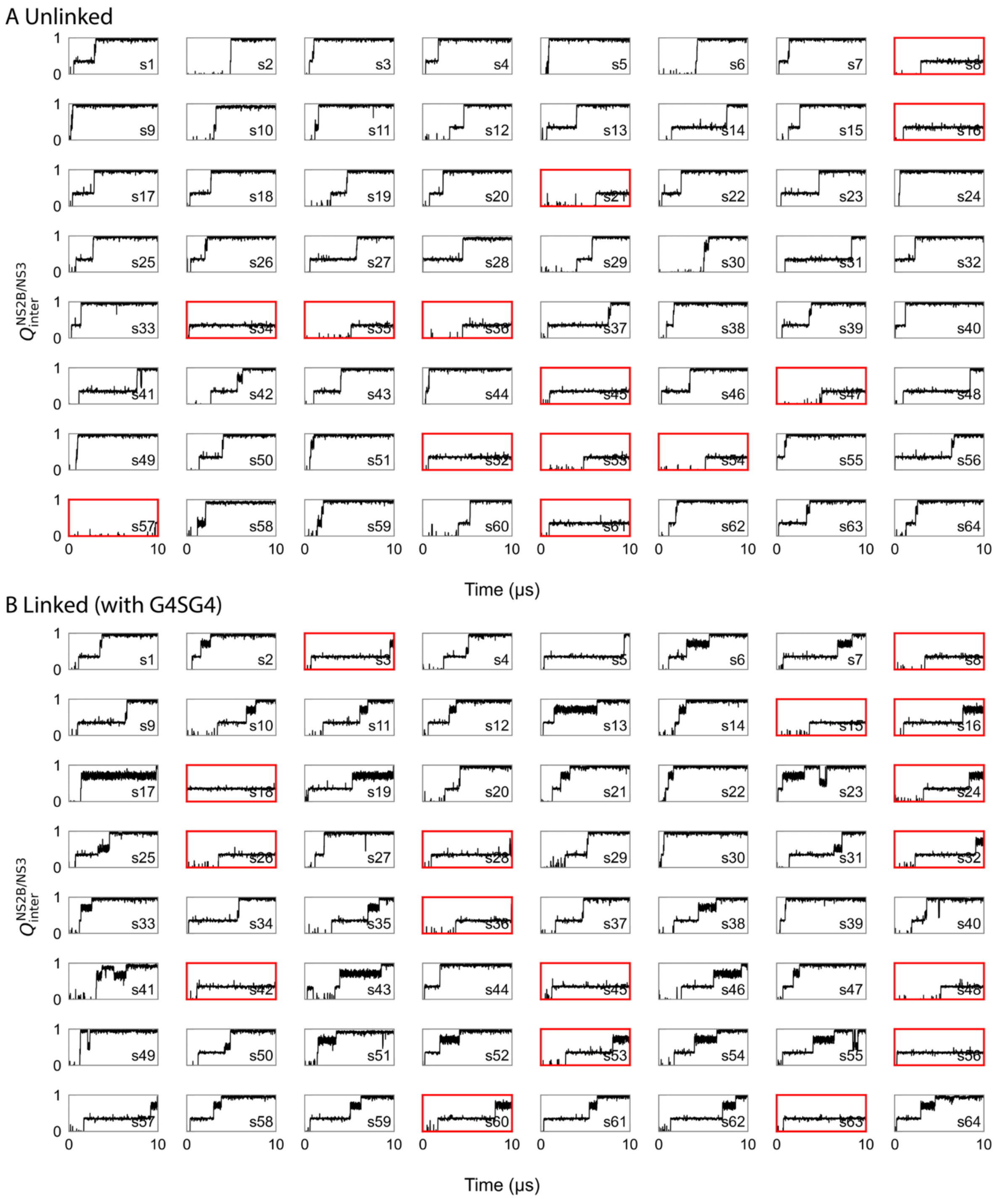


**Supplementary Figure 6. Fraction of native contacts formed between NS2B and NS3 during all 64 folding simulations.** $Q_{inter}^{NS2B,NS3}$ is used as a reference of progress for NS2B/NS3 complex formation. A) The top 64 pannels and B) the bottom 64 panels show traces from simulations of unlinked and linked (with G4SG4) construct, respectively. Eleven unlinked traces (s8, s16, s21, s34, s35, s36, s45, s47, s52, s53, s54, s57, and s61) and seventeen linked traces (s3, s8, s15, s16, s18, s24, s26, s28, s32, s36, s42, s45, s48, s53, s56, s60, and s63) that failed to fold are marked with red frames.

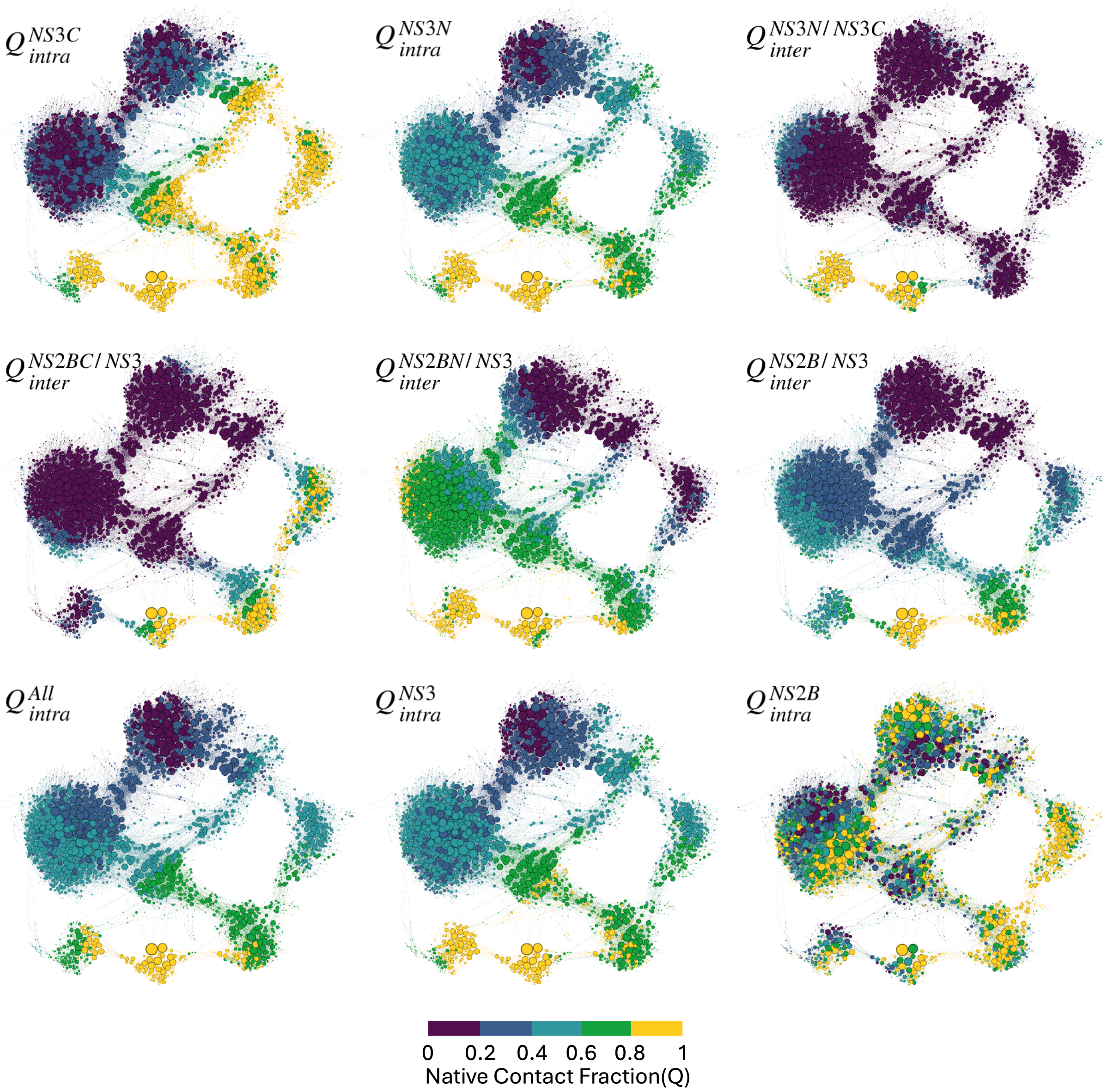


**Supplementary Figure 7. CSNs of folding of the unlinked NS2B/NS3 construct.** Nodes of the same CSN graph after stress minimization are colored using various individual native contact fractions in each panel (see Table 1).

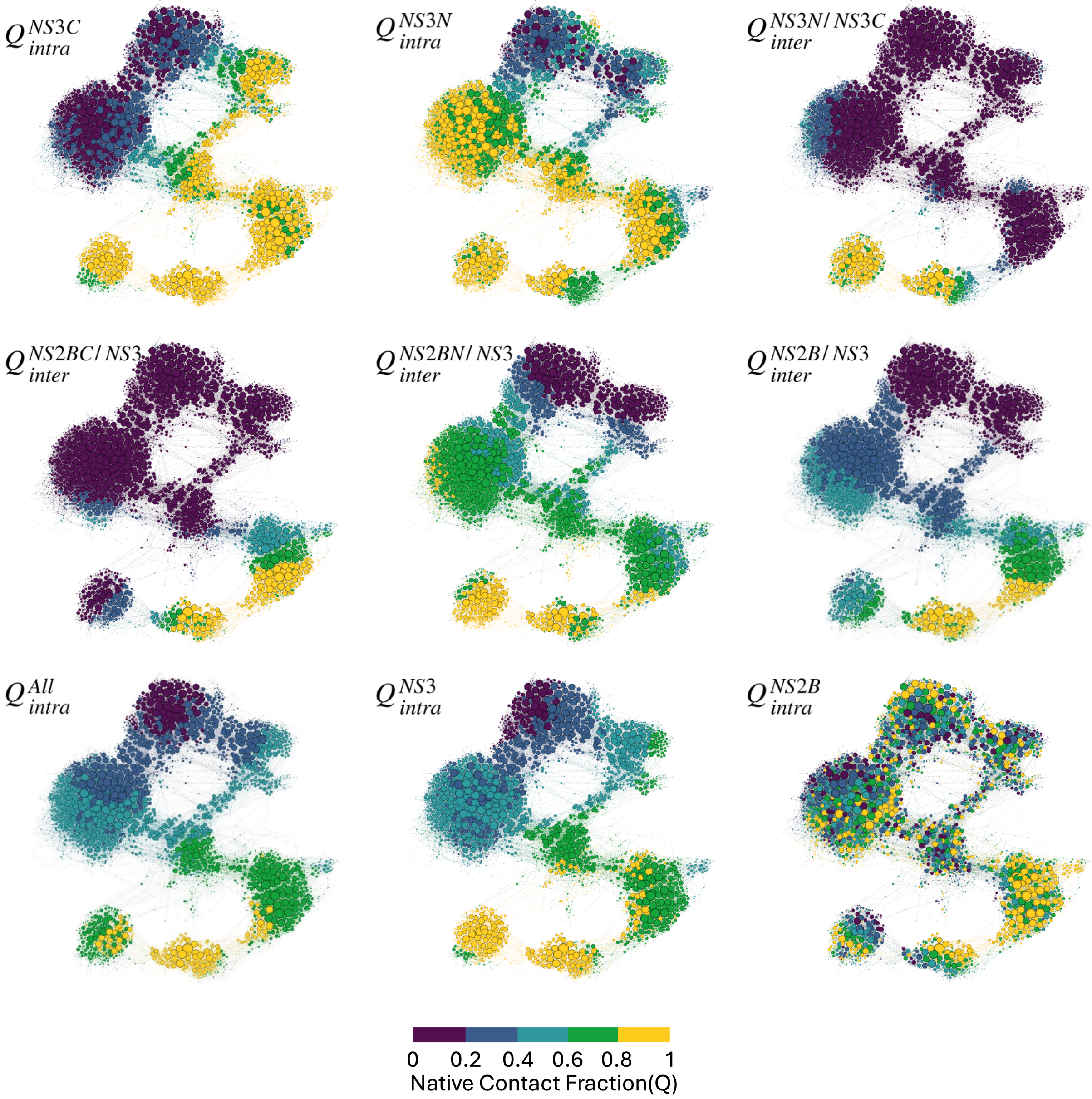


**Supplementary Figure 8. CSNs of folding of the linked NS2B/NS3 construct (with $G_4SG_4$).** Nodes of the same CSN graph after stress minimization are colored using various individual native contact fractions in each panel (see Table 1).

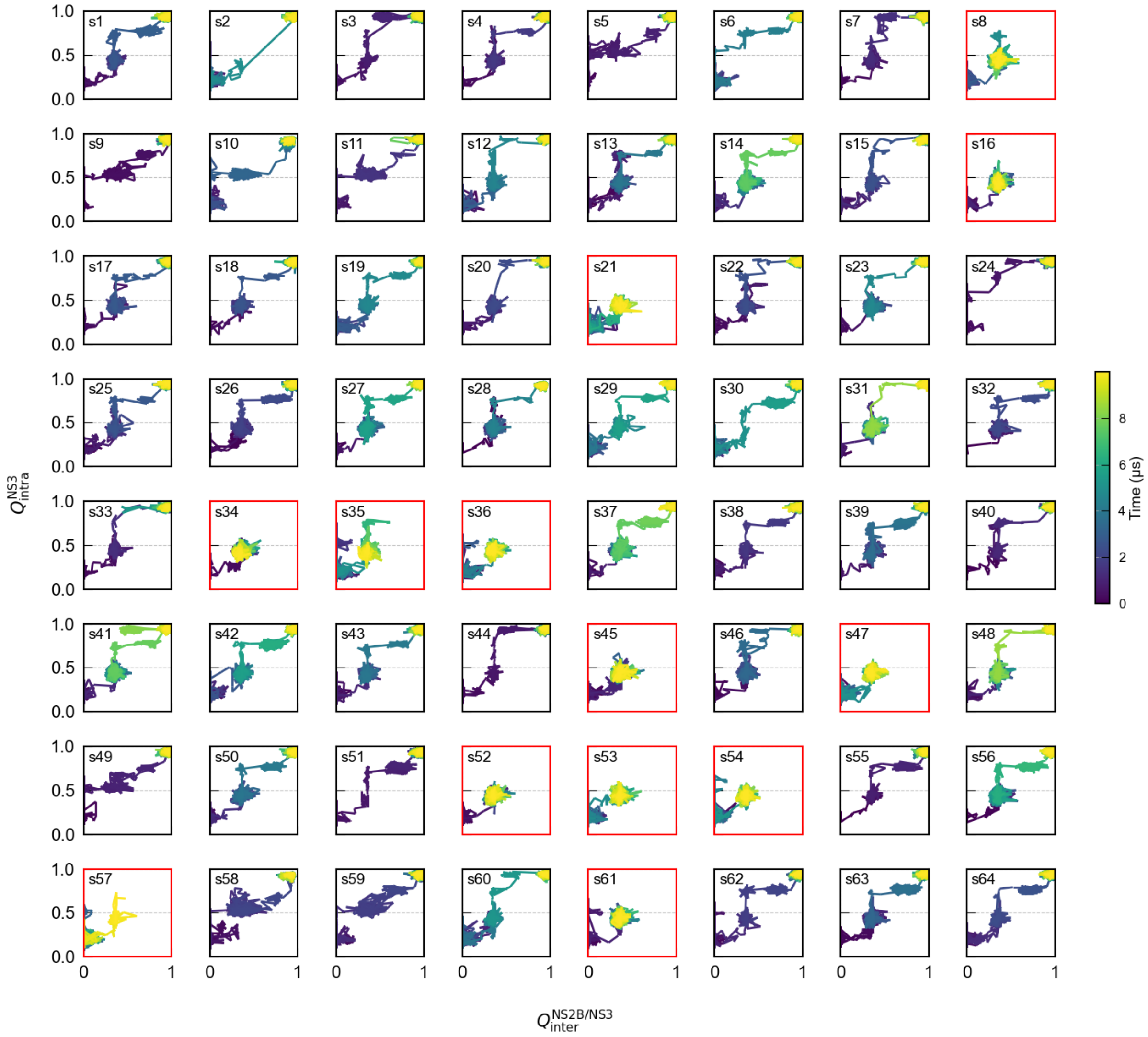


**Supplementary Figure 9. Evolution of native contact fractions $\mathbf{Q_{inter}^{NS2B-NS3}}$ and $\mathbf{Q_{intra}^{NS3}}$ during all 64 folding trajectories of the unlinked NS2B/NS3 construct.** The 13 unlinked trajectories (s8, s16, s21, s34, s35, s36, s45, s47, s52, s53, s54, s57, and s61) that failed to fold are marked with red frames.

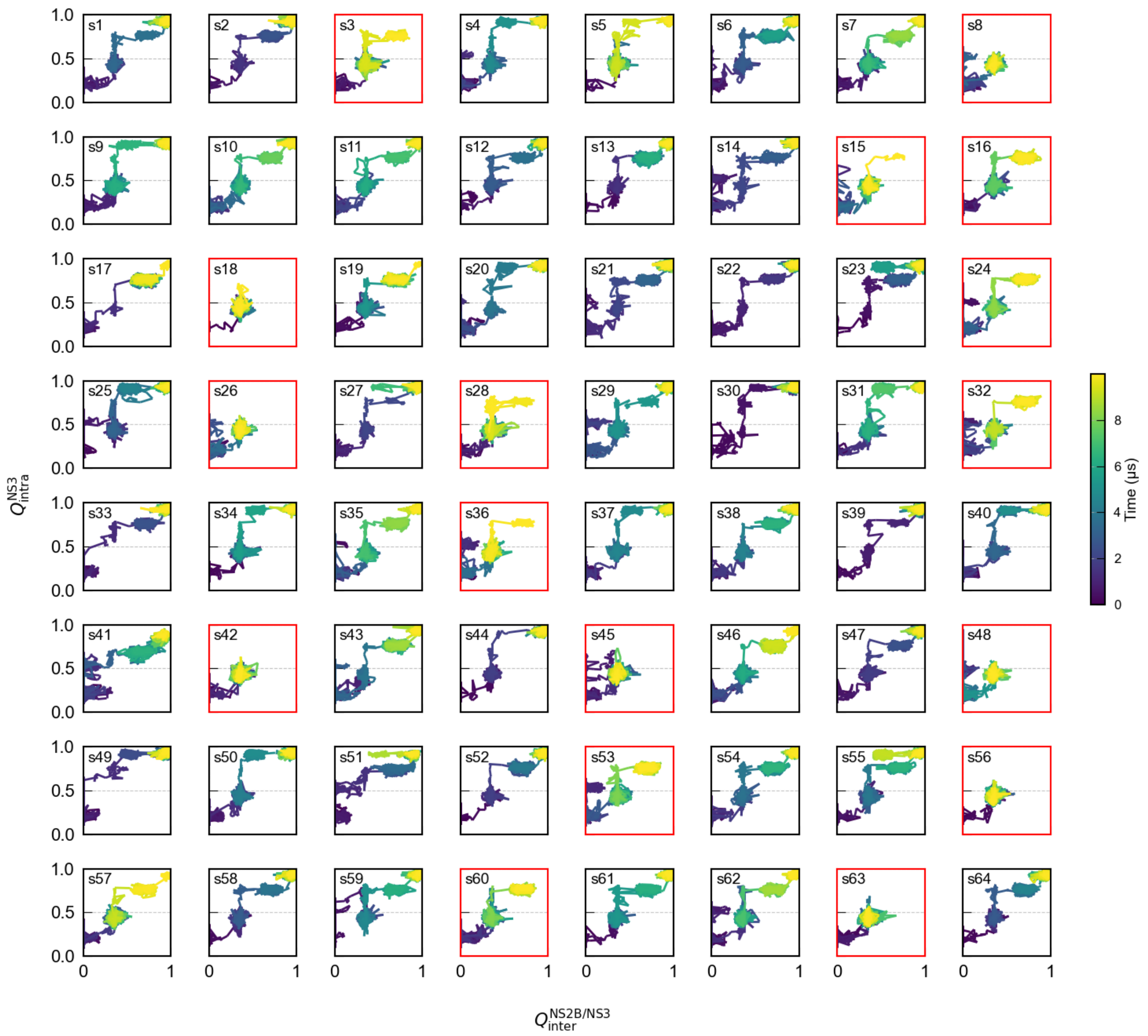


**Supplementary Figure 10. Evolution of native contact fractions $Q_{inter}^{NS2B-NS3}$ and $Q_{intra}^{NS3}$ during all 64 folding trajectories of the linked NS2B/NS3 construct (with $G_4SG_4$).** The 17 linked trajectories (s3, s8, s15, s16, s18, s24, s26, s28, s32, s36, s42, s45, s48, s53, s56, s60, and s63) that failed to fold are marked with red frames.

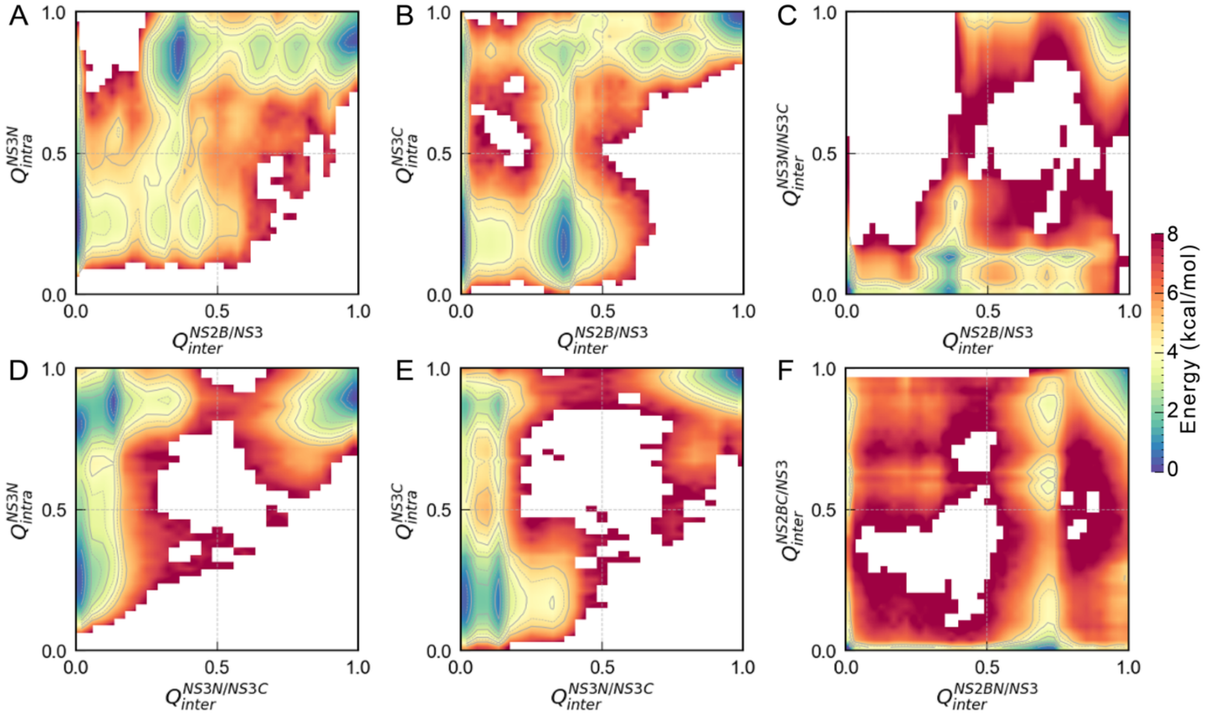


**Supplementary Figure 11. 2D pseudo free energy landscape of folding along various combinations of native contact fractions for the unlinked NS2B/NS3 construct.**

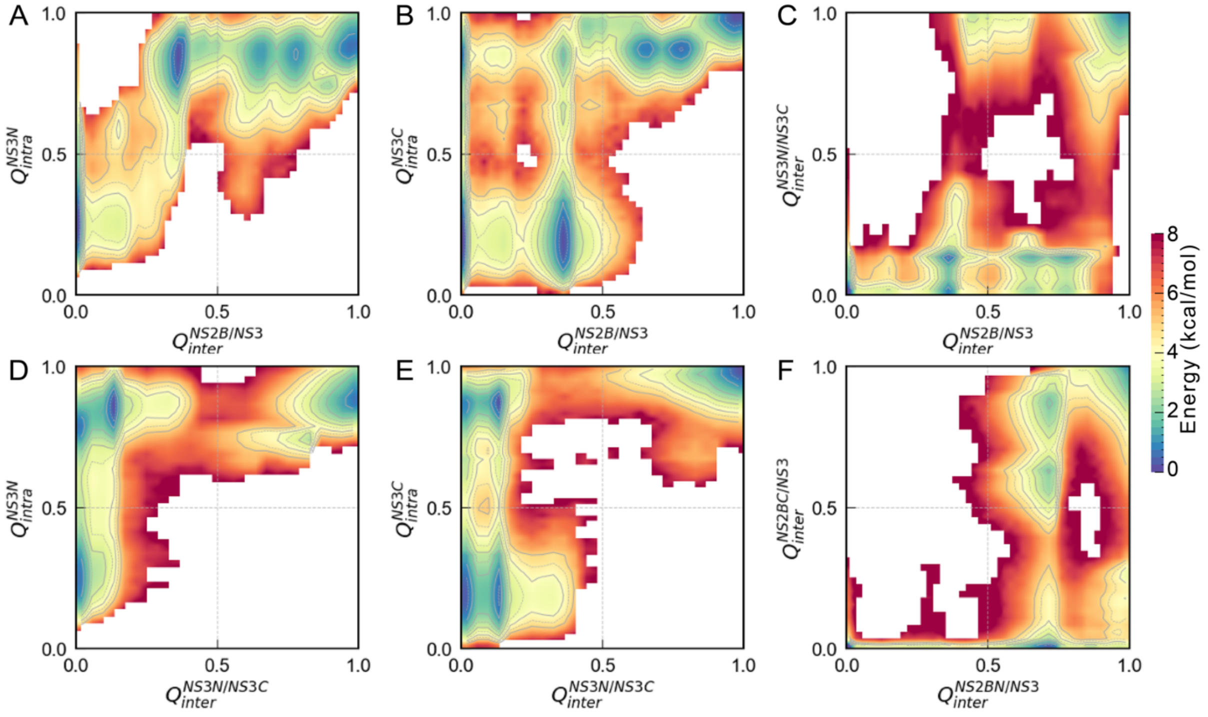


**Supplementary Figure 12. 2D pseudo free energy landscape of folding along various combinations of native contact fractions for the linked NS2B/NS3 construct (with $G_4SG_4$).**

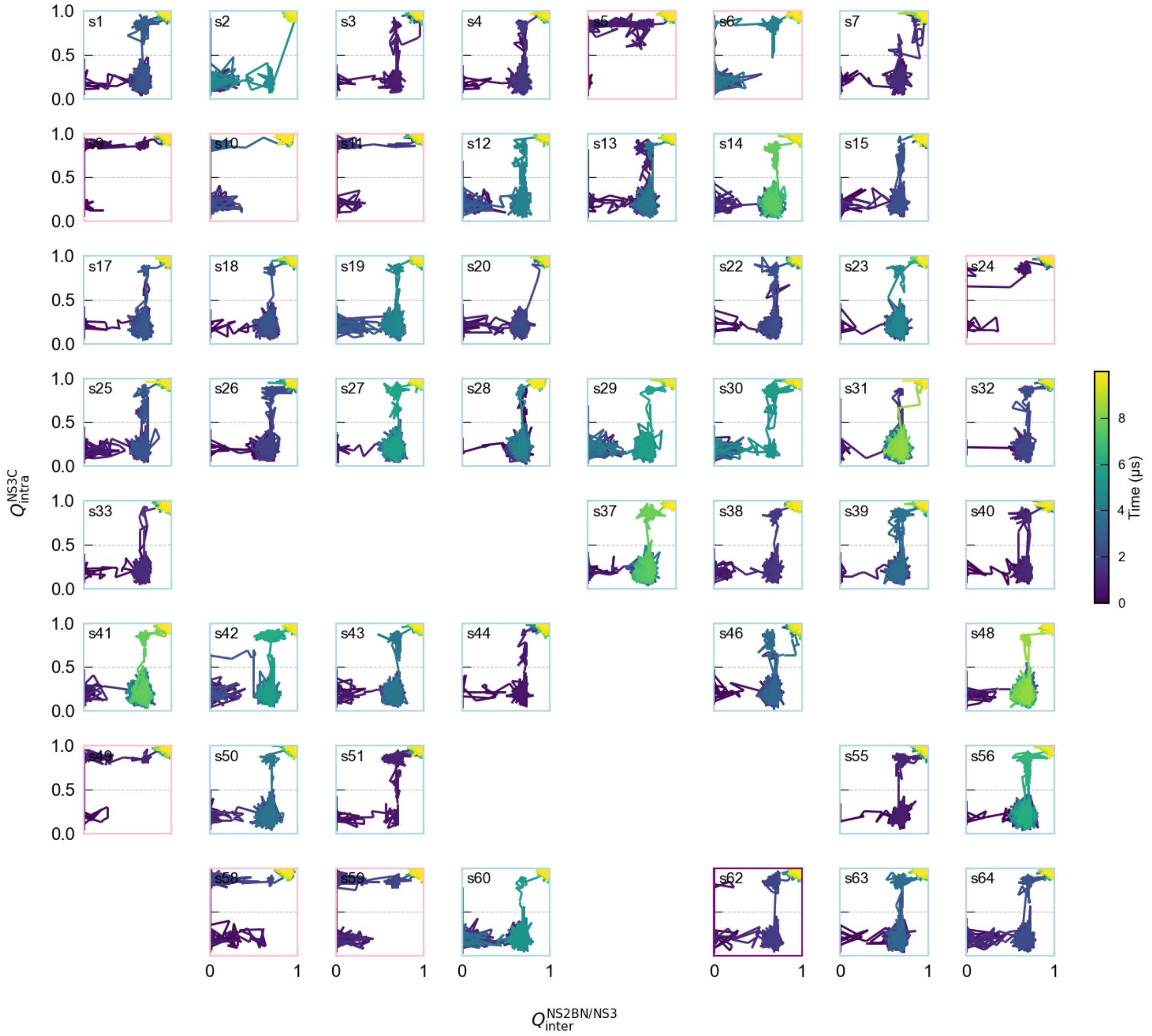


**Supplementary Figure 13. Evolution of native contact fractions $Q_{inter}^{NS2BN-NS3}$ and $Q_{intra}^{NS3C}$ during all 51 successful refolding trajectories of the unlinked NS2B/NS3 construct.** 41 trajectories (s1, s2, s3, s4, s7, s12, s13, s14, s15, s17, s18, s19, s20, s22, s23, s25, s26, s27, s28, s29, s30, s31, s32, s33, s37, s38, s39, s40, s41, s42, s43, s44, s46, s48, s50, s51, s55, s56, s60, s63, and s64) that traversed the I2 state are marked with light blue frames. The 9 trajectories (s5, s6, s9, s10, s11, s24, s49, s58, and s59) that traversed the I1 state are marked with pink frames. The 1 trajectory (s62) that traversed both the I1 and I2 states is marked with purple frames.

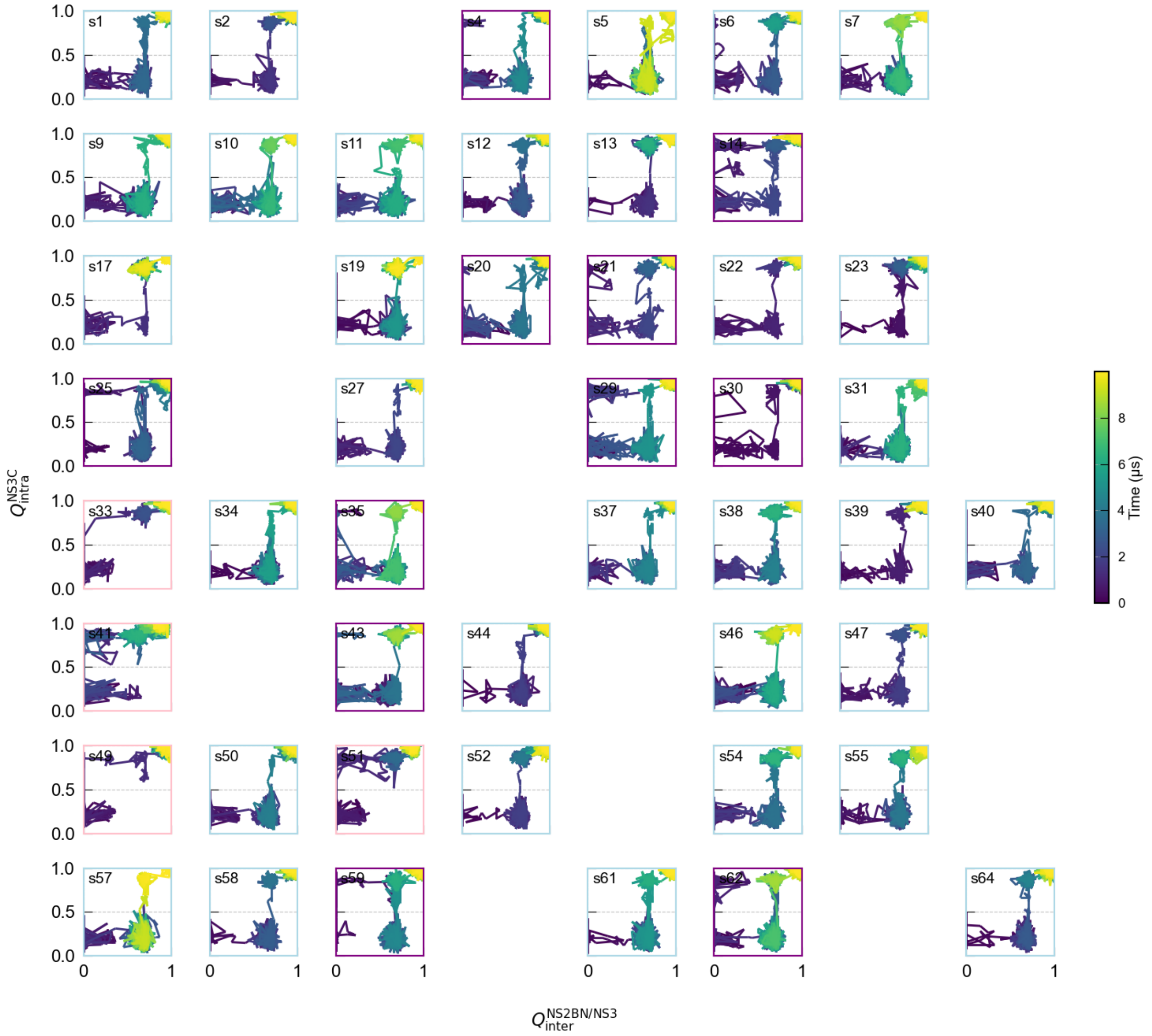


**Supplementary Figure 14. Evolution of native contact fractions $Q_{inter}^{NS2BN-NS3}$ and $Q_{intra}^{NS3C}$ during all 47 successful refolding trajectories of the linked NS2B/NS3 construct (with G4SG4).** The 32 trajectories (s1, s2, s5, s6, s7, s9, s10, s11, s12, s13, s17, s19, s22, s23, s27, s31, s34, s37, s38, s39, s40, s44, s46, s47, s50, s52, s54, s55, s57, s58, s61, and s64) that traversed the I2 state are marked with light blue frames. The 4 trajectories (s33, s41, s49, and s51) that traversed the I1 state are marked with pink frames. The 11 trajectories (s4, s14, s20, s21, s25, s29, s30, s35, s43, s59, and s62) that traversed both the I1 and I2 states are marked with purple frames.

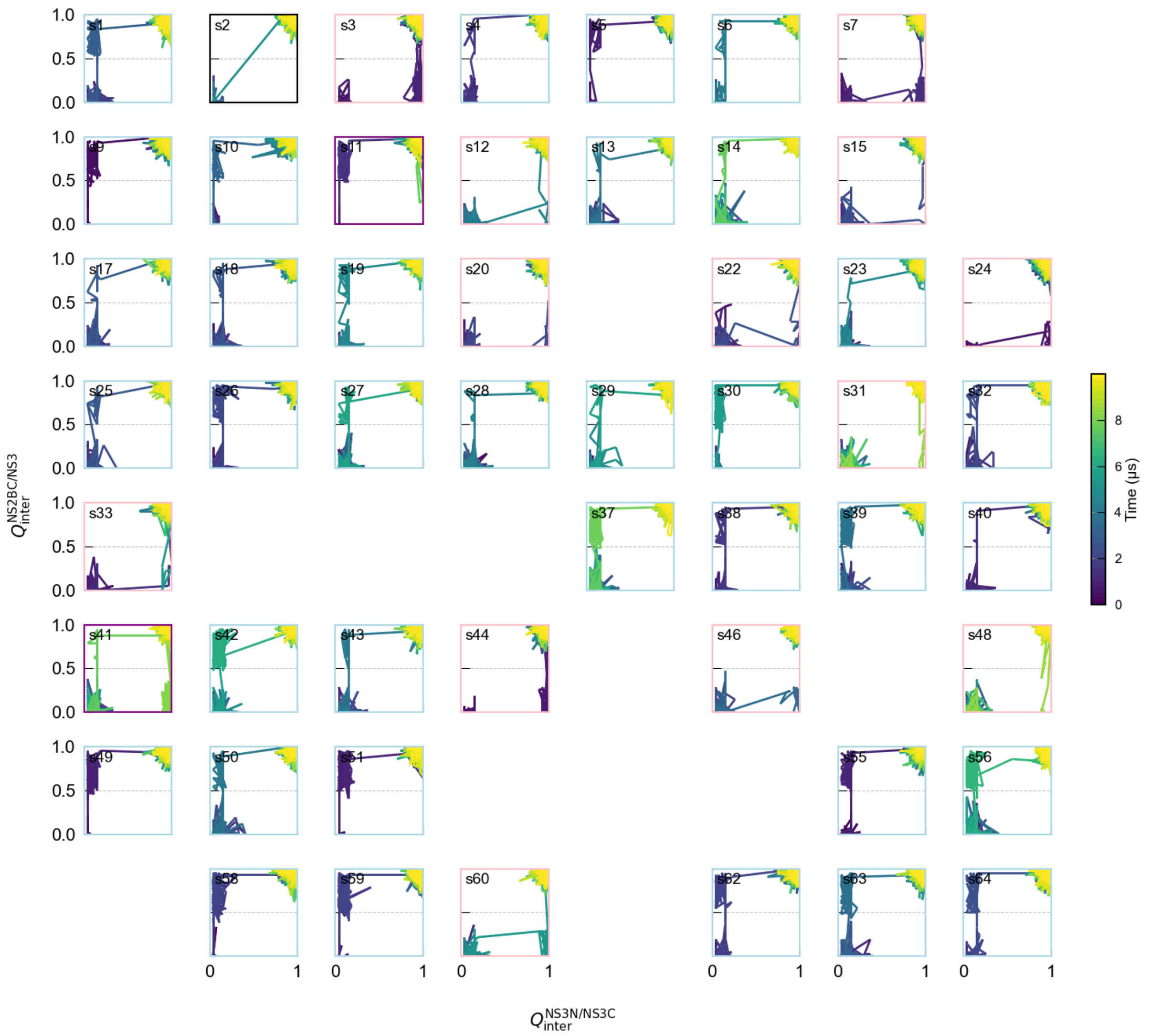


**Supplementary Figure 15. Evolution of native contact fractions $Q_{\text{inter}}^{\text{NS3N−NS3C}}$ and $Q_{\text{inter}}^{\text{NS2BC−NS3}}$ during all 51 successful refolding trajectories of the unlinked NS2B/NS3 construct.** The 35 trajectories (s1, s4, s5, s6, s9, s10, s13, s14, s17, s18, s19, s23, s25, s26, s27, s28, s29, s30, s32, s37, s38, s39, s40, s42, s43, s49, s50, s51, s55, s56, s58, s59, s62, s63, and s64) that traversed the I5 state are marked with light blue frames. The 13 trajectories (s3, s7, s12, s15, s20, s22, s24, s31, s33, s44, s46, s48, and s60) that traversed the I6 state are marked with pink frames. The 2 trajectories (s11 and s41) that traversed both the I5 and I6 states are marked with purple frames.

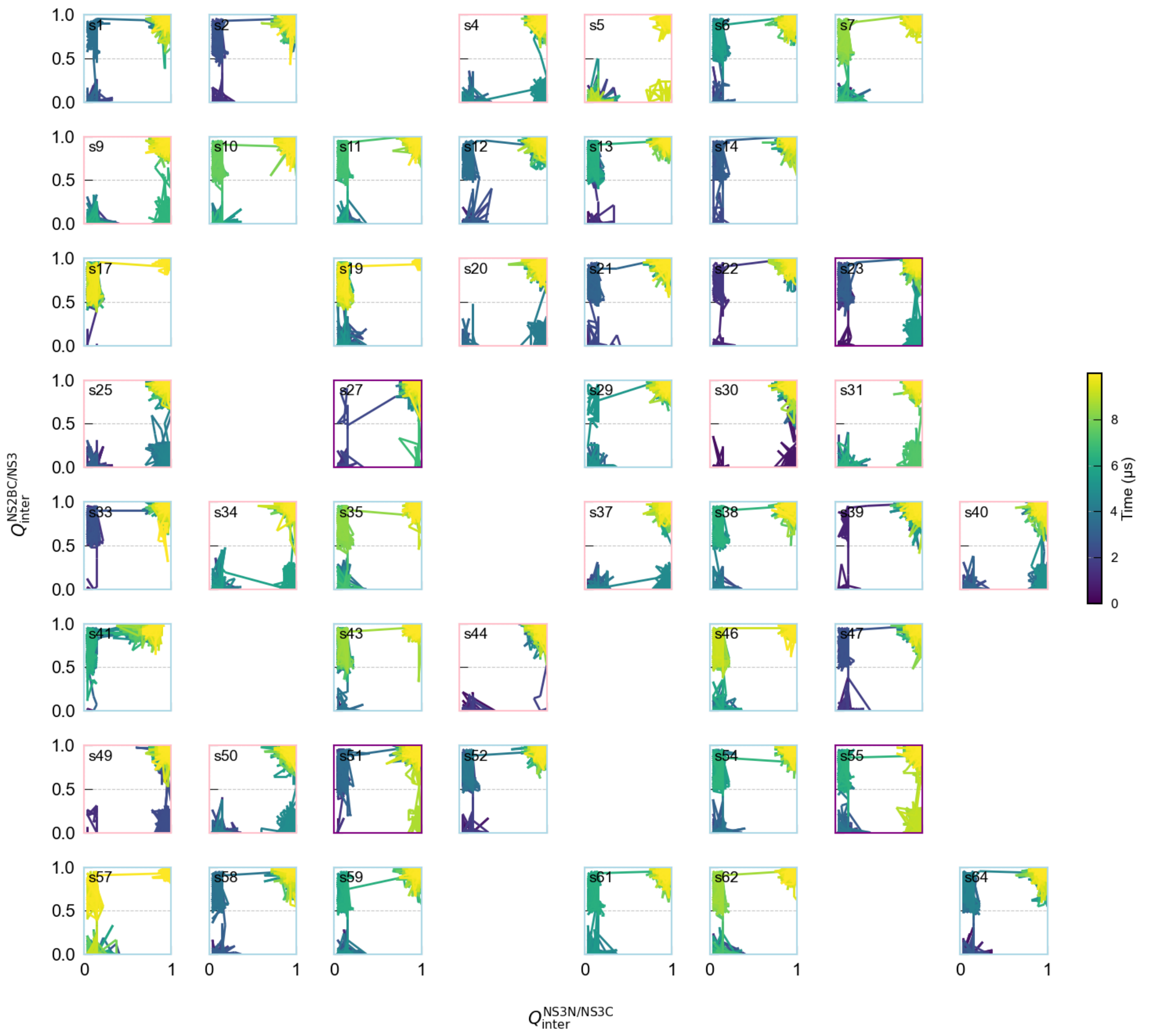


**Supplementary Figure 16. 2D Evolution of native contact fractions $Q_{inter}^{NS3N-NS3C}$ and $Q_{inter}^{NS2BC-NS3}$ during all 47 successful refolding trajectories of the unlinked NS2B/NS3 construct (with G4SG4).** The 30 trajectories (s1, s2, s6, s7, s10, s11, s12, s13, s14, s17, s19, s21, s22, s29, s33, s35, s38, s39, s41, s43, s46, s47, s52, s54, s57, s58, s59, s61, s62, and s64) that traversed the I5 state are marked with light blue frames. The 13 trajectories (s4, s5, s9, s20, s25, s30, s31, s34, s37, s40, s44, s49, and s50) that traversed the I6 state are marked with pink frames. The 4 trajectories (s23, s27, s51, and s55) that traversed both the I5 and I6 states are marked with purple frames.

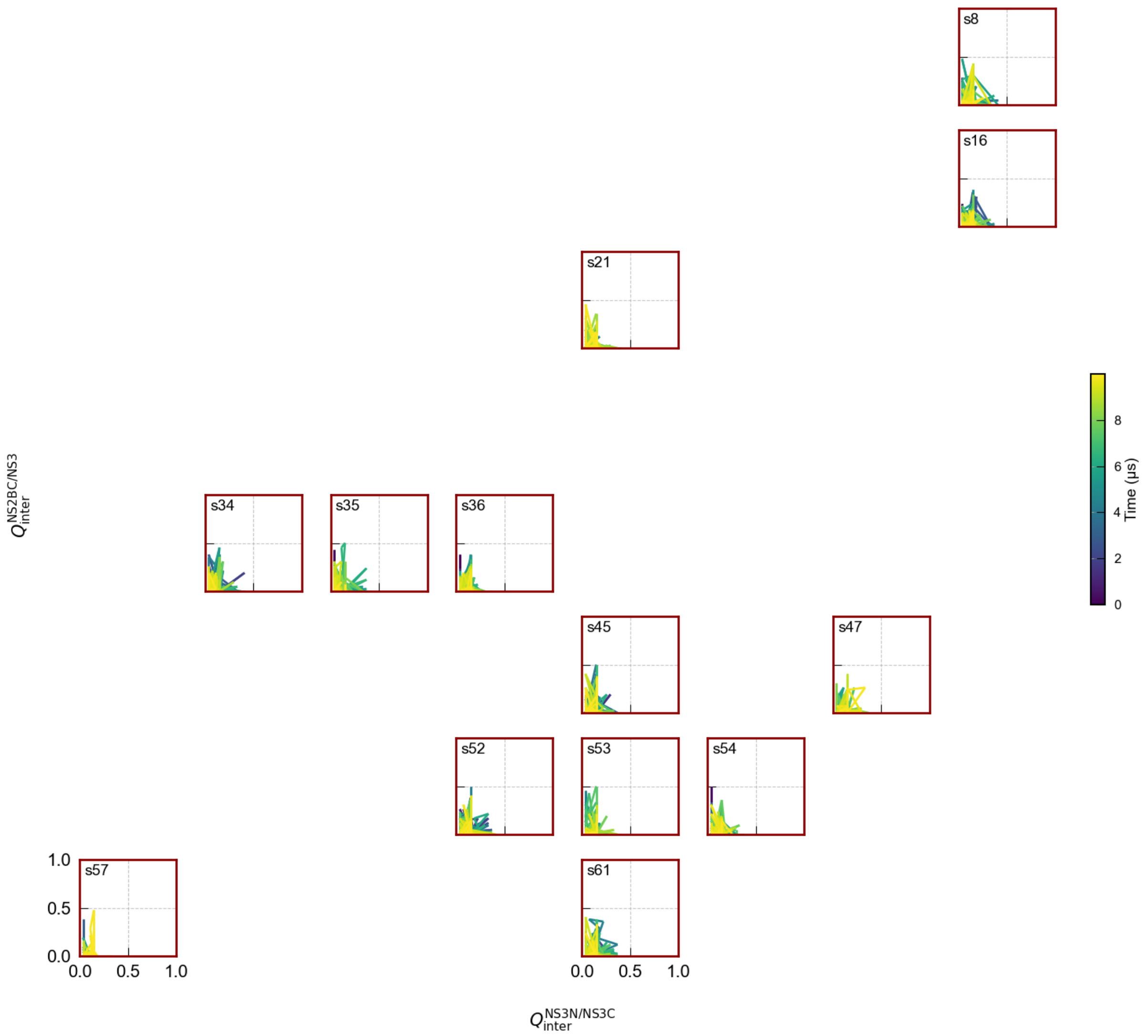


**Supplementary Figure 17. Evolution of native contact fractions $Q_{inter}^{NS3N,NS3C}$ and $Q_{inter}^{NS2BC,NS3}$ during all 13 unsuccessful refolding trajectories of the unlinked NS2B/NS3 construct.** The 13 trajectories (s8, s16, s21, s34, s35, s36, s45, s47, s52, s53, s54, s57, and s61) that failed to fold and became trapped in the I2 state are labeled with dark red frames.

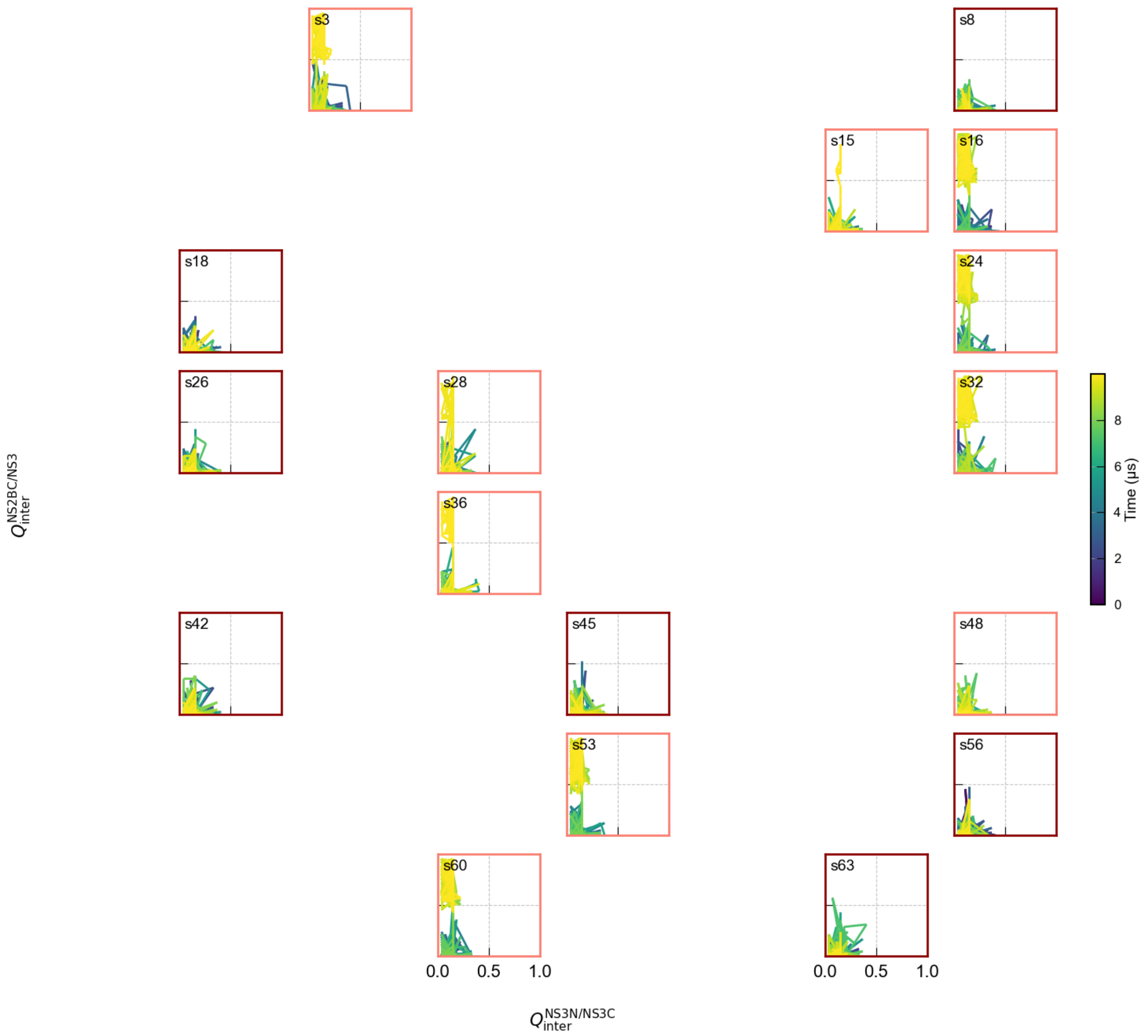


**Supplementary Figure 18. Evolution of native contact fractions $Q_{\mathrm{inter}}^{\mathrm{NS3N-NS3C}}$ and $Q_{\mathrm{inter}}^{\mathrm{NS2BC-NS3}}$ during all 17 unsuccessful refolding trajectories of the linked NS2B/NS3 construct (with G4SG4).** The 7 trajectories (s8, s18, s26, s42, s45, s56, and s63) that failed to fold and became trapped in the I5 state are shown with dark red frames. The 10 trajectories (s3, s15, s16, s24, s28, s32, s36, s48, s53, and s60) that failed to fold and became trapped in the I5 state are shown with salmon-colored frames.

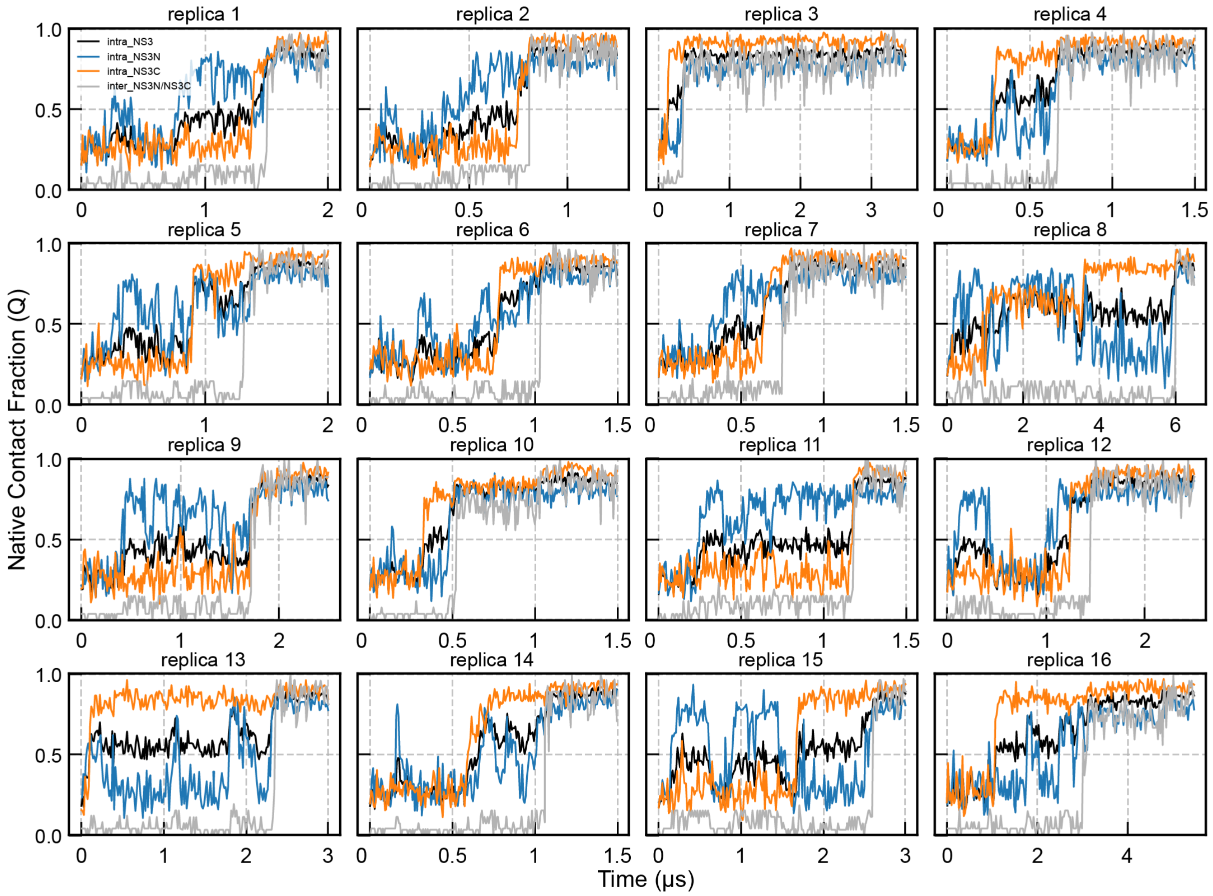


**Supplementary Figure 19. Evolution of various native contact fractions during folding simulations of NS3 at 300 K.** Traces for $Q^{NS3}_{intra}$ ; $Q^{NS3N}_{intra}$ ; $Q^{NS3C}_{intra}$ ; $Q^{NS3N-NS3C}_{inter}$ are colored black, blue, orange and gray, and labeled as “intra_NS3”, “intra_NS3N”, “intra_NS3” and “intra_NS3N/NS3C” respectively. The generation of initial unfolded configurations and the followed 16 folding simulations were performed using the same protocol as used for NS2B/NS3 construct (see Methods).

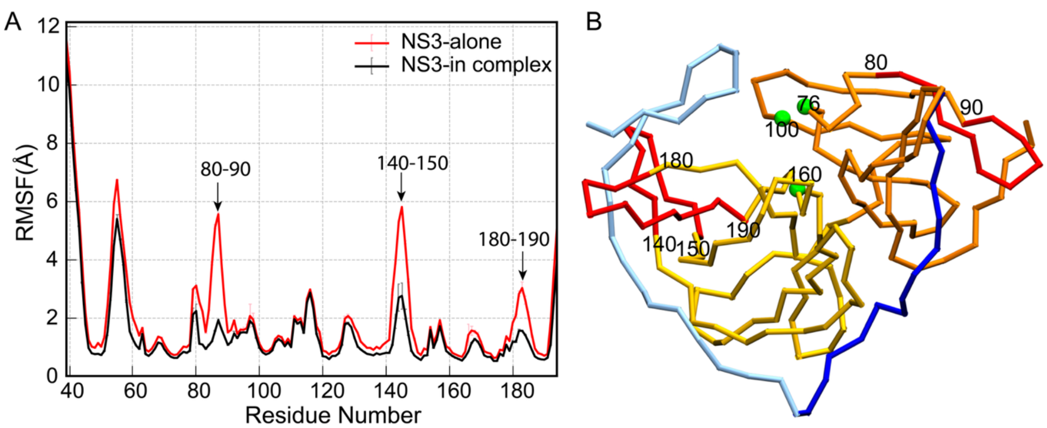


**Supplementary Figure 20. Structure and dynamics of NS3 by itself and in complex with NS2B. A)** RMSF profiles of NS3 derived from Go model simulation of NS3 in complex with NS2B or simulated as NS3 alone with three loops with the largest differences marked. **B**) Trace representation of the NS2B/NS3 complex with the three loops colored in red and the catalytic triad residues marked with green spheres.
